# An ancient clade of *Penelope*-like retroelements with permuted domains is present in the green lineage and protists, and dominates many invertebrate genomes

**DOI:** 10.1101/2021.04.23.441226

**Authors:** Rory J. Craig, Irina A. Yushenova, Fernando Rodriguez, Irina R. Arkhipova

**Affiliations:** Institute of Evolutionary Biology, University of Edinburgh, Edinburgh, UK; Josephine Bay Paul Center for Comparative Molecular Biology and Evolution, Marine Biological Laboratory, Woods Hole, MA, USA

**Author notes:** These authors contributed equally to this work (Rory J. Craig, Irina A. Yushenova).

**Keywords:** transposable elements, retrotransposons, reverse transcriptase, GIY-YIG endonuclease, selenoproteins, microsatellites

## Abstract

*Penelope*-like elements (PLEs) are an enigmatic clade of retroelements whose reverse transcriptases (RTs) share a most recent common ancestor with telomerase RTs. The single ORF of canonical EN+ PLEs encodes RT and a C-terminal GIY-YIG endonuclease (EN) that enables intrachromosomal integration, while EN–PLEs lack endonuclease and are generally restricted to chromosome termini. EN+ PLEs have only been found in animals, except for one case of horizontal transfer to conifers, while EN–PLEs occur in several kingdoms. Here we report a new, deep-branching PLE clade with a permuted domain order, whereby an N-terminal GIY-YIG endonuclease is linked to a C-terminal RT by a short domain with a characteristic Zn-finger-like motif. These N-terminal EN+ PLEs share a structural organization, including pseudo-LTRs and complex tandem/inverted insertions, with canonical EN+ PLEs from *Penelope/Poseidon*, *Neptune* and *Nematis* clades, and show insertion bias for microsatellites, but lack hammerhead ribozyme motifs. However, their phylogenetic distribution is much broader. The *Naiad* clade is found in numerous invertebrate phyla, where they can reach tens of thousands of copies per genome. *Naiads* in spiders and clams independently evolved to encode selenoproteins. *Chlamys*, which lack the CCHH motif universal to PLE endonucleases, occur in green algae, spike mosses (targeting ribosomal DNA) and the slime mold *Physarum*. Unlike canonical PLEs, RTs of N-terminal EN+ PLEs contain the insertion-in-fingers domain, strengthening the link between PLEs and telomerases. Additionally, we describe *Hydra*, a novel metazoan C-terminal EN+ clade. Overall, we conclude that PLE diversity, distribution and abundance is comparable to non-LTR and LTR-retrotransposons.

## INTRODUCTION

Transposable elements (TEs) are characterized by their intrinsic ability to move within and between genomes. In eukaryotes, TEs contribute not only to structural organization of chromosomes and variation in genome size, but also to genetic and epigenetic regulation of numerous cellular processes (Wells and Feschotte 2020). TEs are traditionally divided into two classes, based on the presence (class I, retrotransposons) or absence (class II, DNA transposons) of an RNA intermediate in the transposition cycle. Retrotransposons, in turn, are divided into subclasses based on the presence or absence of long terminal repeats (LTRs): LTR-retrotransposons are framed by direct repeats, phylogenetically close DIRS elements by split inverted repeats, non-LTR retrotransposons lack terminal repeats, and *Penelope*-like elements (PLEs) have a special kind of repeats called pseudo-LTRs (pLTRs), which may be in direct or inverted orientation. Repeat formation in each subclass is associated with the combined action of phylogenetically distinct clades of reverse transcriptase (RT) domain fused to different types of endonuclease/phosphotransferase (EN) domains: DDE-type integrases (IN) or tyrosine recombinases (YR) in LTR-retrotransposons; restriction enzyme-like (REL) or apurinic /apyrimidinic (AP) endonucleases in non-LTR retrotransposons; and GIY-YIG endonucleases in PLEs (Arkhipova 2017). The fusion of EN to RT is typically C-terminal, with the exception of an N-terminal EN in *copia*-like LTR-retrotransposons and in AP-containing non-LTR retrotransposons. The concerted action of RT and EN that combines cleavage and joining of DNA strands with cDNA synthesis during retrotransposition results in characteristic terminal structures that define the boundaries of new insertions.

The GIY-YIG EN domain typically associated with PLEs may have its evolutionary origins in bacterial group I introns, which are not retroelements (Stoddard 2014). The group I intron-encoded homing ENs are characterized by long recognition sequences, and act essentially as monomeric nickases, cleaving DNA on one strand at a time. The relatively short GIY-YIG cleavage module (~70 aa) is often tethered to additional DNA-binding domains for target recognition (Derbyshire, et al. 1997; Van Roey, et al. 2002).

In eukaryotic PLEs, the activity of the recombinant GIY-YIG EN has been studied *in vitro* for *Penelope* elements of *Drosophila virilis*, where it displayed several properties expected from homology to prokaryotic enzymes, such as functional catalytic residues, nicking activity producing a free 3’-OH for RT priming, and moderate target preferences (Pyatkov, et al. 2004). Variable distance between first-strand cleavage of DNA during target-primed reverse transcription (TPRT) and second-strand cleavage upon TPRT completion dictates the variable length of the target-site duplication (TSD), which is observed at the integration site. Phylogenetically, PLE ENs form a distinct cluster within a large GIY-YIG nuclease superfamily, where diverse homing ENs occupy a central position (Dunin-Horkawicz, et al. 2006). PLE ENs are distinguished from those of homing ENs by the presence of a highly conserved CCHH Zn-finger motif, where the two cysteines are located directly between the GIY and YIG motifs (Arkhipova 2006).

Phylogenetic history of the longer RT domain is much more informative and reveals a sister relationship between PLEs and telomerase RTs (TERTs), which add G-rich telomeric repeats to extend eukaryotic chromosome ends (Arkhipova, et al. 2003). All described PLEs form two major groups: endonuclease-deficient (EN–) PLEs, retroelements found in several eukaryotic kingdoms at or near telomeres, and endonuclease-containing (EN+) PLEs, which harbor a C-terminal GIY-YIG EN enabling retrotransposition throughout the genome (Fig. 1) (Gladyshev and Arkhipova 2007). Three large EN+ PLE clades have been named *Penelope/Poseidon*, *Neptune* and *Nematis*, the latter two being characterized by the presence of an additional conserved Zn-finger motif in the linker between RT and EN (Arkhipova 2006). Two EN–RT clades, *Athena* and *Coprina*, lack the EN domain entirely, but display a unique ability to attach to exposed G-rich telomeric repeat overhangs, assisted by stretches of reverse-complement telomeric repeats combined with adjacent hammerhead ribozyme motifs (HHR) (Gladyshev and Arkhipova 2007; Arkhipova, et al. 2017). Despite the ancient origin of PLEs predating their divergence from TERTs, which are pan-eukaryotic, the phylogenetic distribution of EN+ PLEs has so far been restricted to animals, with one exception of documented horizontal transfer to conifers (Lin, et al. 2016). Here we report the discovery of a novel deep-branching EN+ PLE clade, where the GIY-YIG EN is unexpectedly positioned N-terminally to the RT. A clade of these elements present in animals, termed *Naiad*, contains the GIY-YIG domain bearing the characteristic Zn-fingers found in canonical EN+ PLEs, while a second group, termed *Chlamys*, are present in green algae, spike mosses and the slime mold *Physarum*, and lack both EN Zn-finger motifs. These results uncover hitherto unknown PLE diversity, which spans all eukaryotic kingdoms, testifying to their ancient origins. We also report that *Naiads* from species as diverse as spiders and clams can code for selenoproteins, which have not previously been described in any TEs.

**Fig. 1.**
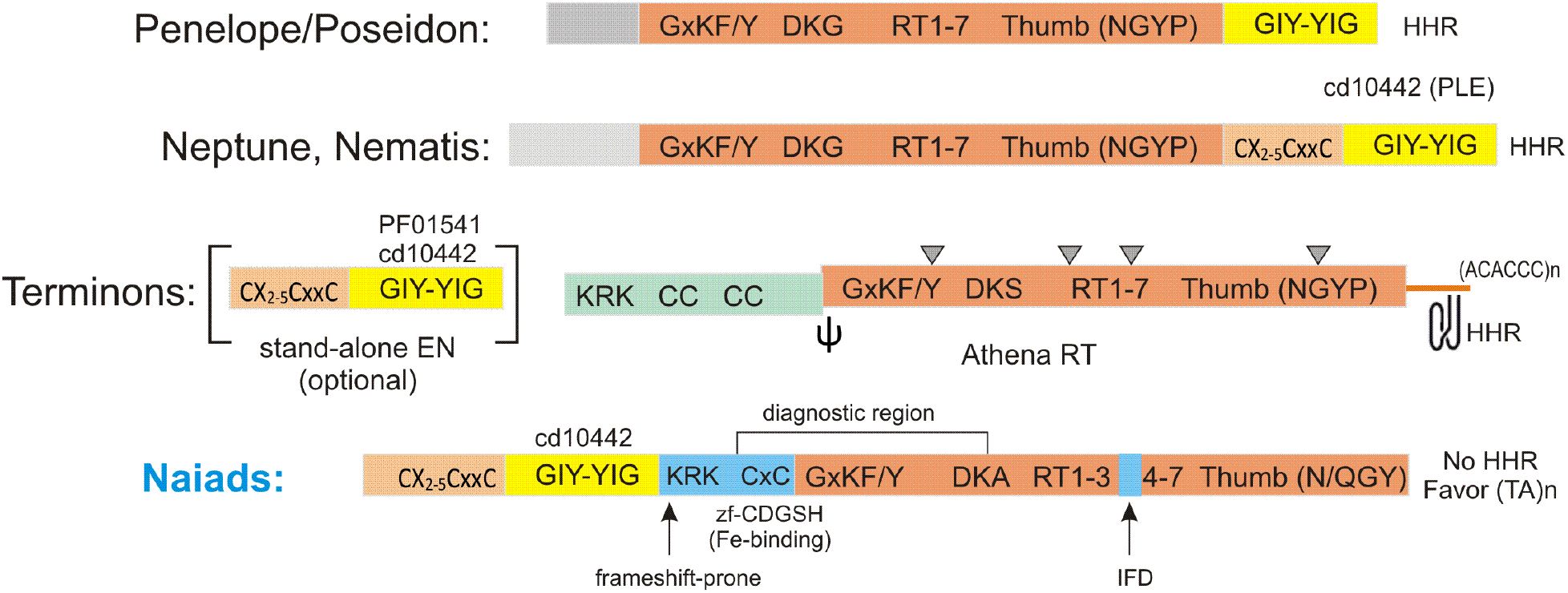
Domain architecture of the major PLE types found in animals. Domains are colored as follows: RT (peach), GIY-YIG (yellow), ORF1 (pink) often separated by a frameshift and a pseudoknot (Ψ), Zn-fingers (sand), and N-terminal domains with no characteristic motifs (gray). The organization of *Coprina* elements from fungi, protists and plants is similar to *Athena*. *Naiad*-specific domains are in blue. CC, coiled-coil; IFD, insertion in fingers domain; HHR, hammerhead ribozyme motif (also present in pLTRs of canonical EN+ PLEs); KRK, nuclear localization signal; (ACACCC)_n_, short stretches of reverse-complement telomeric repeats in EN-deficient PLEs. Conserved introns are denoted by triangles. Also shown are the most conserved amino acid motifs and the highest-scoring PFAM/CD domain matches. Not to scale.

## RESULTS

### Novel PLEs with N-terminal location of the GIY-YIG endonuclease domain

While cataloguing PLEs in several recently sequenced genomes, such as the acanthocephalan (*Pomphorhynchus laevis*) and a bdelloid rotifer (*Didymodactylos carnosus*), as well as a darwinulid ostracod (*Darwinula stevensoni*) (Mauer, et al. 2020; Nowell, et al. 2021; Schön, et al. 2021), we noticed the absence of the GIY-YIG domain at the C-terminus of several PLEs, which is typically indicative of EN–PLEs. In these cases, however, extending the 5’-end of the frequently truncated PLE copies revealed a conserved N-terminal GIY-YIG EN domain, typically 220-275 aa in length. A high degree of 5’-truncation apparently precluded earlier identification of this novel type of PLEs. For instance, Repbase, a comprehensive database of eukaryotic TEs (Bao, et al. 2015), contains two PLEs consistently appearing as top RT matches to the novel PLEs, yet having no N-terminal EN domain (*Penelope-2_CGi* from the Pacific oyster *Crassostrea gigas* and *Penelope-1_EuTe* from the Texas clam shrimp *Eulimnadia texana*). We extended the 767-aa *Penelope-2_CGi_1p* consensus in the 5’-direction and compared it with two sibling species, *Crassostrea virginica* and especially *Saccostrea glomerata*, where this element is mostly intact, revealing an N-terminal GIY-YIG domain which brings the total ORF length up to 876 aa in *C. gigas* (still 5’-truncated) and to 1024 aa in *S. glomerata*.

We then conducted an extensive database search for representatives of this previously undescribed type of PLEs in sequenced genomes, relying primarily on the N-terminal position of the GIY-YIG domain and several characteristic motifs (see below) to discriminate between novel and canonical PLEs (Fig. 1). Our search revealed a surprising diversity of hosts from eight animal phyla, including ctenophores, cnidarians, rotifers, nematodes, arthropods, mollusks, hemichordates, and vertebrates (fish). Additionally, about a dozen hits on short contigs were annotated as bacterial, however upon closer inspection these were discarded as eukaryotic contaminants from metagenomic assemblies with an incorrect taxonomic assignment (Arkhipova 2020). Out of 36 animal host species, most were aquatic (26), 6 were parasitic, and only 4 were free-living terrestrial species (2 spiders and 2 nematodes). We therefore chose the name *Naiad* for this newly discovered type of PLEs.

### Structural characteristics of Naiad elements

Structurally, *Naiad* insertions exhibit most of the previously known characteristic features of PLEs (Evgen’ev and Arkhipova 2005; Arkhipova 2006). Insertions show a high degree of 5’-truncation and are often organized into partial tandems, so that a full-length copy would be preceded by a partially truncated copy, forming a pLTR. Often, there is also an inverted 5’-truncated copy found immediately adjacent at the 5’-end, leading to formation of inverted repeats flanking the entire insertion unit (Fig. 2). Such complex structures of insertions often lead to problems in WGS assembly, especially short read-based. To further complicate boundary recognition, a 30-40 bp extension (“tail”) is usually found at either end of the insertion unit, most likely resulting from EN-mediated resolution of the transposition intermediate. However, a notable difference between *Naiads* and canonical PLEs is the absence of any detectable hammerhead ribozyme motifs (HHR), which are typically located within pLTRs (Cervera and De la Peña 2014; Arkhipova, et al. 2017). Ignoring any tandemly inserted sequence, the main body of full-length *Naiad* copies are generally 3.4 - 4.4 kb.

**Fig. 2.**
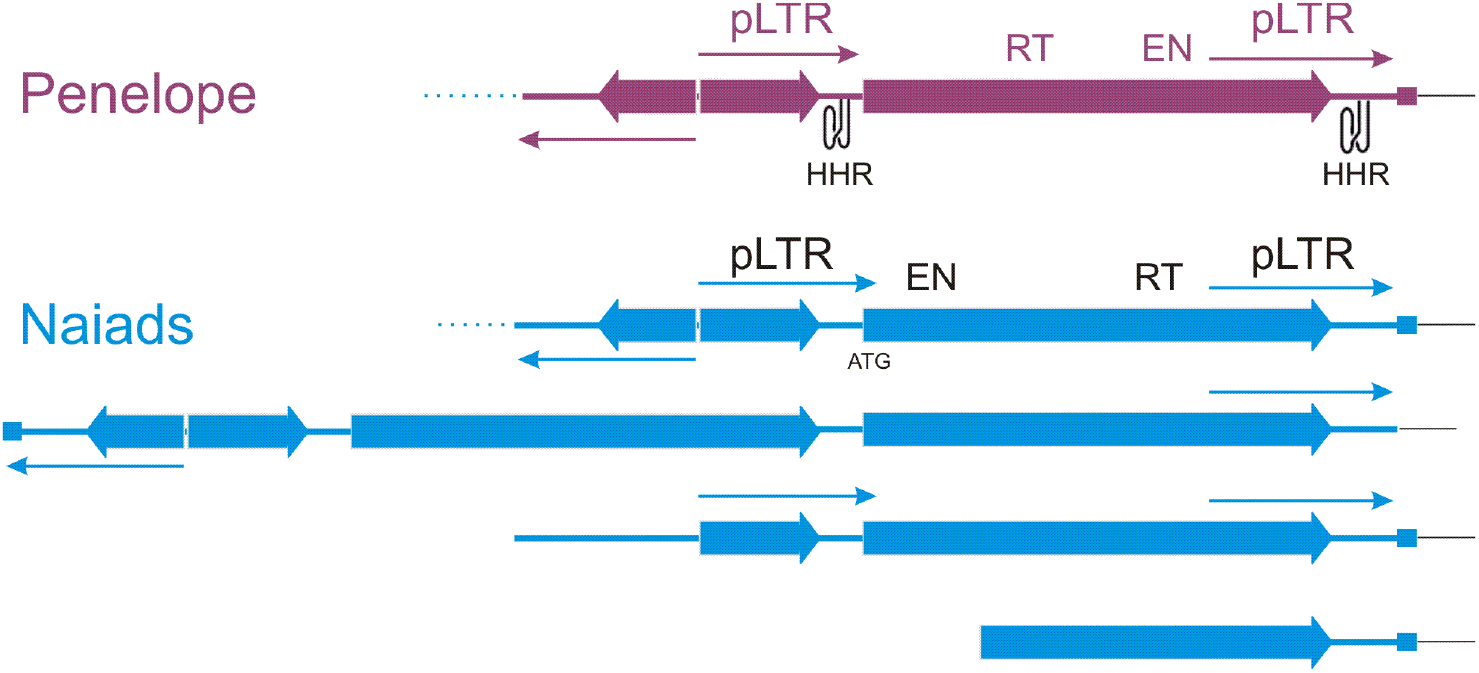
Structural arrangements of PLE copies. Shown is the typical arrangement of two or more ORFs in partial tandems, forming pseudo-LTRs (pLTRs) denoted by arrows. An inverted 5’-truncated copy is often found adjacent to the upstream pLTR, forming an inverted-repeat structure. A small square denotes a 30-40 bp extension (“tail”) that is usually present only on one end of the insertion. HHR, hammerhead ribozyme motif. For *Penelope*, only the most typical structure is shown, but all other variants also observed.

Sequence conservation of the RT domain is strong enough to retrieve RTs of canonical PLEs in a BLAST search, thus it is practical to rely on several diagnostic regions, such as the CxC motif (showing weak homology to zf-CDGSH Fe-binding Zn-fingers) and the DKG motif (Arkhipova 2006), which in *Naiads* is modified to DKA (Fig. 1). In the core RT, the region between RT3(A) and RT4(B) is ~20 aa longer than in other PLEs and corresponds in position to the IFD (insertion in the fingers domain) of TERTs (Lingner, et al. 1997; Lue, et al. 2003) (Fig. S1). Interestingly, the IFD is missing from *Naiads* in chelicerates (spiders and the horseshoe crab) and *D. stevensoni*, which resemble canonical PLEs in this region. Finally, between RT and the upstream EN domain there is usually a large KR-rich block harboring a nuclear localization signal (Fig. 3B). This block, which is rich in adenines, is particularly prone to frameshift mutations resulting in detachment of the EN domain from RT and its eventual loss. Such mutations apparently prevented earlier recognition of the EN domain in *C. gigas* and *E. texana* PLEs from Repbase (Bao, et al. 2015).

**Fig. 3.**
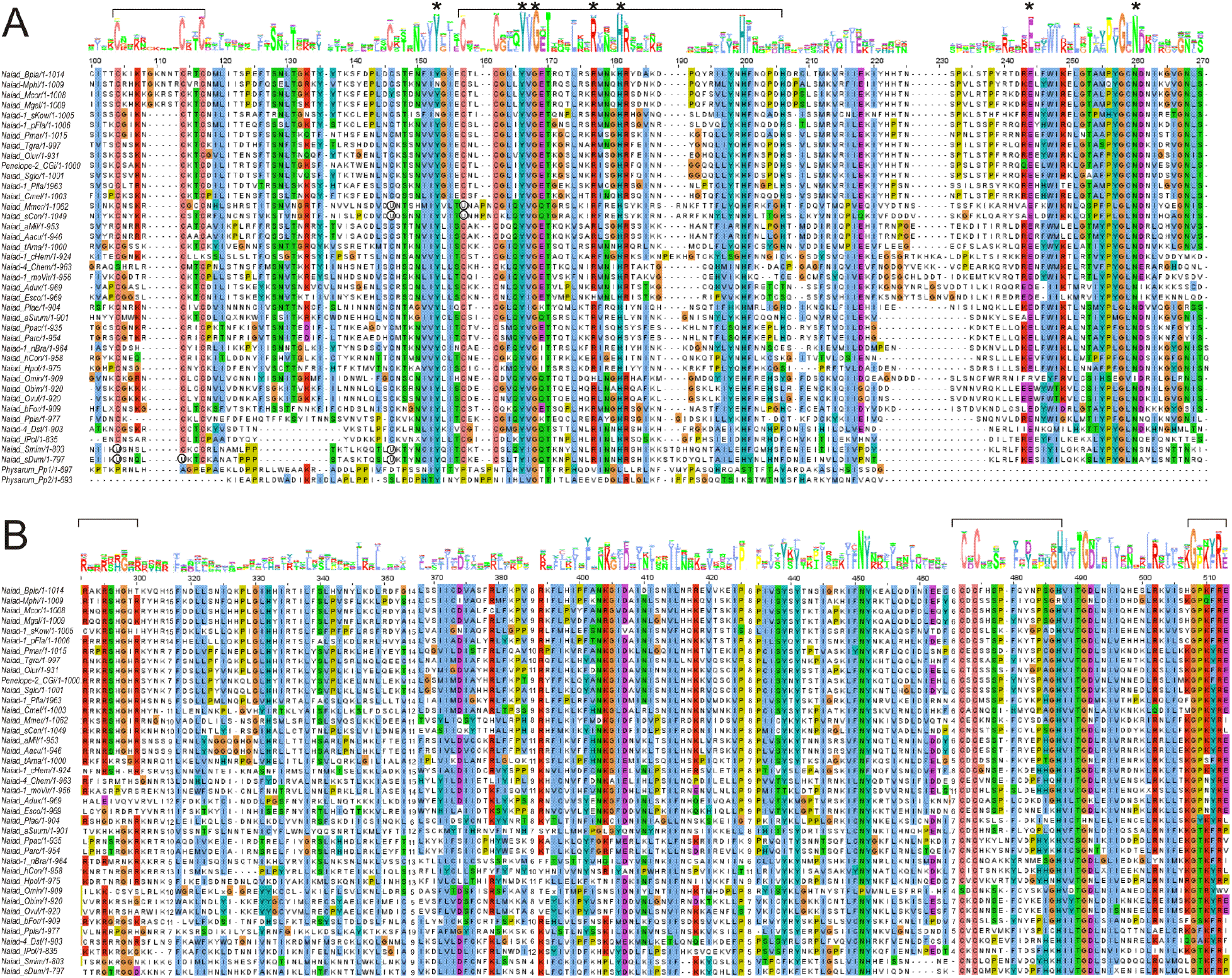
Multiple sequence alignments of conserved domains characteristic for Naiads. (**A**) The GIY-YIG EN domain. The Zn-finger-like motifs are demarcated by square brackets; catalytic residues are denoted by asterisks; residues corresponding to selenocysteines (U) are circled. The position of the second His in the CCHH motif is variable. (**B**) The conserved diagnostic region between the EN domain and the GxKF/Y motif present in other PLEs. The KR-rich, CxC and GxKF/Y motifs are marked by square brackets. Alignments were visualized in Jalview and the sequence logos were created by AlignmentViewer (Waterhouse, et al. 2009; Gabler, et al. 2020), using the Clustal2 coloring scheme. The sequence order corresponds to that in Fig. 5.

The EN domain in *Naiads* displays most similarity to the GIY-YIG EN of other PLEs (cd10442), especially those in the *Neptune* and *Nematis* clades which harbor an additional conserved CX_2-5_CxxC Zn-finger-like motif (lengthened by 5-aa insert in mussels) upstream from the GIY-YIG motif (Fig. 1, 3A, S2). Perhaps it may facilitate recognition of (TA)_n_ microsatellite sequences, which often serve as preferred targets for *Naiad* insertion. Its designation as a Zn-finger is tentative, as it shows variably non-significant matches to ZnF_NFX, ZnF_A20, ZnF_TAZ, ZnF_U1 or RING fingers in SMART database searches (Letunic, et al. 2020). The CCHH Zn-finger-like motif with two cysteines inside the GIY-YIG core, characteristic of all canonical EN+ PLEs (Arkhipova 2006), is also present, and the catalytic domain beyond the GIY-YIG core is well conserved and includes the R, H, E and N residues implicated in catalysis (Van Roey, et al. 2002). Thus, despite the permuted arrangement of the RT and EN domains, *Naiads* share the peculiarities of structural organization with other PLEs, indicating that their retrotransposition likely proceeds through a similar mechanism.

### Naiads can reach exceptionally high copy numbers

While PLEs in animal genomes are typically outnumbered by non-LTR and LTR retrotransposons, we noticed that *Naiads* can be particularly successful in certain genomes in comparison to known PLE types. For example, inspection of TE landscape divergence profiles in the acanthocephalan *P. laevis* (Fig. 4A) shows that PLE families are responsible for 8.9% of the genome, of which *Naiad-1* occupies 7.8%. The remaining 12 *Neptune* and 4 *Penelope/Poseidon* families combined occupy only 1.1% of the genome (Fig. 4B).

**Fig. 4.**
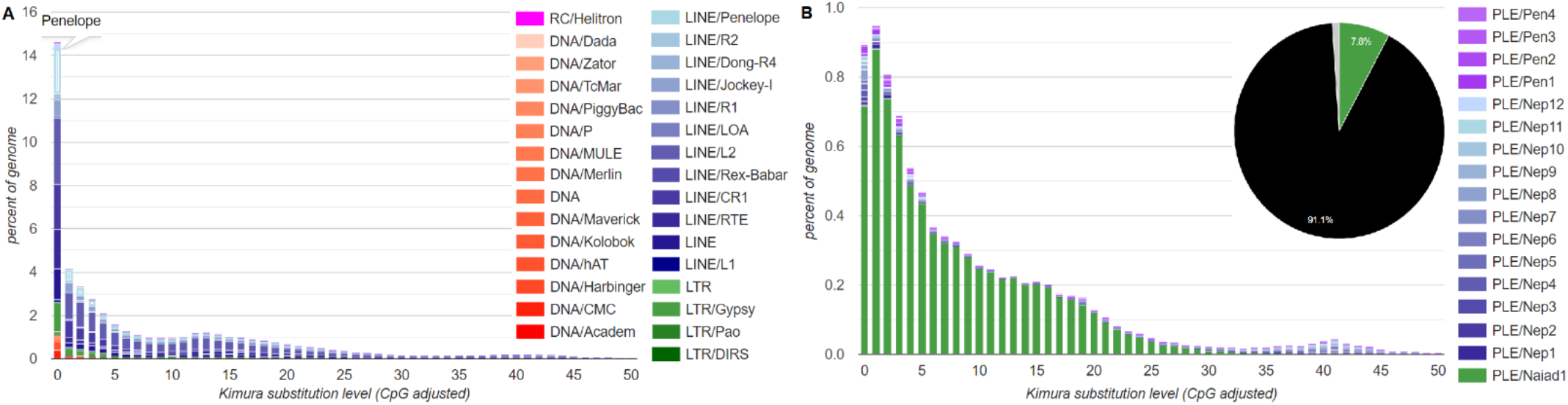
Landscape divergence plots showing TE activity over time and genome occupancy in the acanthocephalan *Pomphorhynchus laevis*. (**A**) All TEs, with PLEs in light blue; (**B**) PLEs only, subdivided by families, with Naiad1_Plae family shown in green on the divergence plot and on the inserted pie chart.

We estimated copy numbers in each host species by querying each WGS assembly with the corresponding *Naiad* consensus sequence and counting the number of 3’-ends at least 80 bp in length. This approach avoids counting multiple fragments in lower-quality assemblies. Among hosts, significant variation in *Naiad* copy number can be observed, even between closely related species (Fig. 5). Copy numbers mostly reflect activity levels: some *Naiads* are apparently intact and are still successfully amplifying, while in other species they have been inactivated a long time ago and required numerous ORF corrections to yield an intact consensus. Surprisingly, several marine invertebrates, such as oysters, clams and crabs, harbor tens of thousands of *Naiad* copies, with nearly 37,000 in the blue crab *Paralithodes platypus*. The lack of HHR motifs obviously has not hampered the proliferative capacity of *Naiads*, as they can outnumber canonical HHR-bearing PLE families in the same species by several orders of magnitude, as in *P. laevis* (Fig. 4B).

**Fig. 5.**
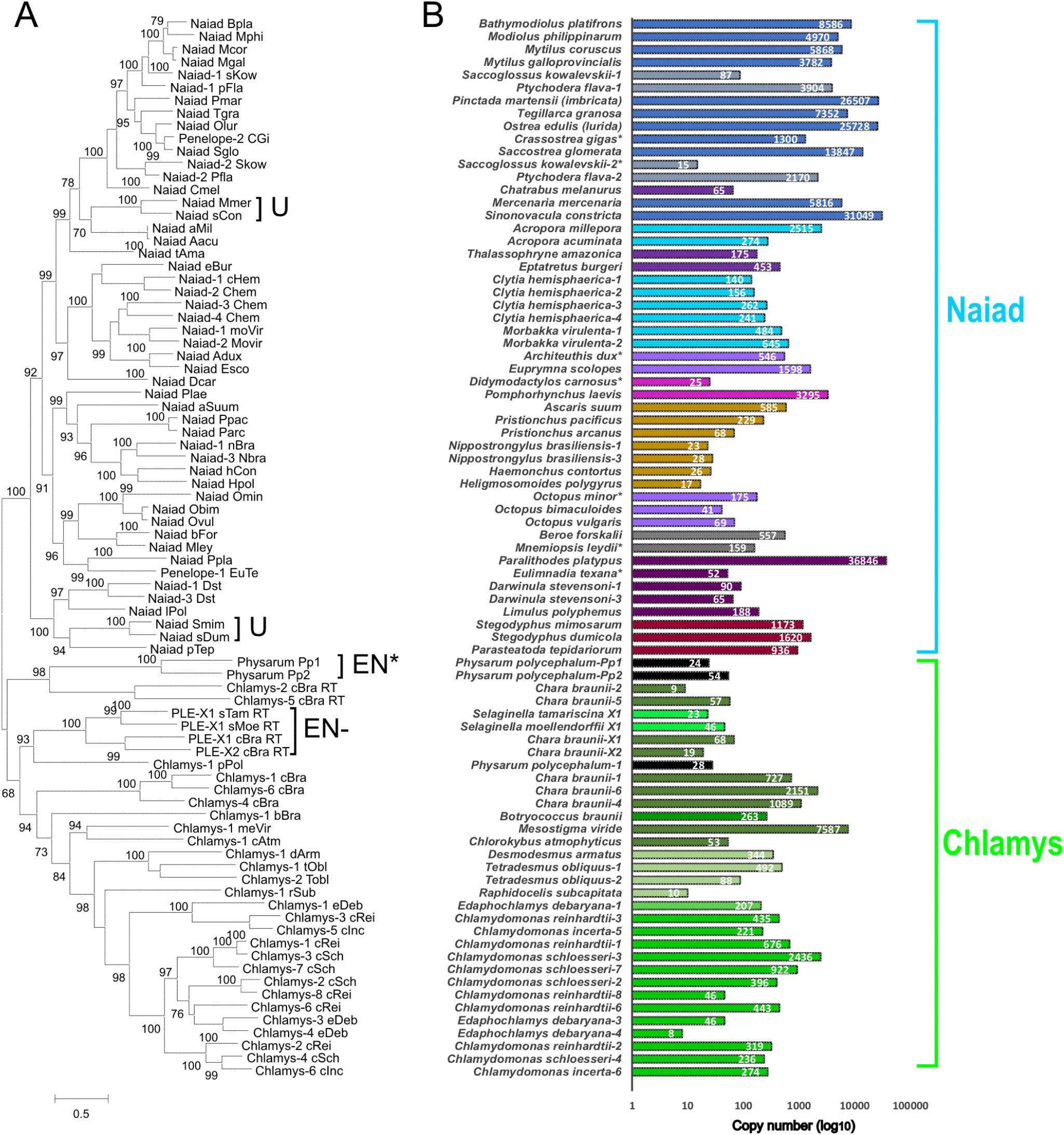
Phylogenetic relationships between different *Naiad* and *Chlamys* families and copy number counts for each family. (**A**) The maximum likelihood phylogram shows branch support from 1000 ultrafast bootstrap replications. Families harboring selenocysteines are denoted by U, families lacking the EN domain by EN-, and families with EN remnants by EN*. Scale bar, aa substitutions per site. (**B**) The copy number chart displays counts for each family on a log scale. Similar colors denote similar taxonomic affiliations. Asterisks mark truncated or interrupted ORFs which are presumably non-functional.

It is also evident that the *Naiad* phylogeny does not necessarily parallel that of host species. While some species, such as *Clytia hemisphaerica* or *D. stevensoni*, have experienced substantial within-species *Naiad* diversification, harboring four families each, others, such as hemichordates (*Saccoglossus kowalevskii*, *Ptychodera flava*), or cephalopods (*Architeuthis dux, Euprymna scolopes*, and *Octopus spp*.), harbor families belonging to different *Naiad* lineages (Fig. 5A). The fish *Naiads* (from *Chatrabus melanurus, Thalassophryne amazonica* and *Eptatretus burgeri*) do not form a monophyletic clade, while the nematode or arthropod *Naiads* do (Fig. 5A). The overall distribution pattern is suggestive of vertical inheritance punctuated by occasional horizontal transfer events and multiple losses.

### Naiad selenoproteins in clams and spiders

The ORFs of four *Naiads*, from two clams (*Sinonovacula constricta* and *Mercenaria mercenaria*) and two spiders (*Stegodyphus dumicola* and *Stegodyphus mimosarum*), each contained either three (*Naiad_Smim* and *Naiad_Mmer*) or four (*Naiad_sDum* and *Naiad_sCon*) in-frame UGA codons. Except for one UGA codon in *Naiad_sCon*, all UGA codons corresponded to highly conserved cysteines in the protein sequences of other *Naiads* (Fig. 3). In all families, UGA codons corresponded to the cysteine preceding the GIY-YIG motif, and the cysteine eight aa downstream of the DKA motif (not shown). In spiders, UGA codons corresponded to either the first (*Naiad_Smim*) or both the first and second (*Naiad_sDum*) cysteines in the CX_2-5_CxxC Zn-finger, while in clams UGA codons corresponded to the first cysteine in the CCHH Zn-finger. The single remaining UGA codon in *Naiad_sCon* corresponded to an aa in RT6(D) that was not strongly constrained.

Given the correspondence between the in-frame UGA codons and conserved cysteines, we hypothesized that the ORFs of these *Naiads* may encode selenoproteins, in which UGA is recoded from stop to selenocysteine (Sec). Recoding is achieved through a *cis*-acting selenoprotein insertion sequence (SECIS), a Sec-specific tRNA and additional *trans*-acting proteins (Berry, et al. 1991; Tujebajeva, et al. 2000). In eukaryotes, SECIS elements are located in the 3’ UTRs of selenoprotein mRNAs (Low and Berry 1996). Using SECISearch3 (Mariotti, et al. 2013) to query each consensus sequence, we identified “grade A” (i.e., the highest confidence) type I SECIS elements in all four of the families (Fig. S3A). Except for *Naiad-Mmer*, the predicted SECIS elements were located immediately downstream of the inferred UAA or UAG stop codons (1 – 21 bp downstream, Fig. S3B), presumably placing the SECIS elements within the 3’ UTRs of each family. In *Naiad-Mmer*, the SECIS overlapped the first non-UGA stop codon, however there was a UGA codon 7 bp upstream of the SECIS. The recoding of UGA is position dependent (Turanov, et al. 2013) and a UGA codon in such close proximity to the SECIS is not expected to efficiently encode Sec (Wen, et al. 1998), suggesting that this UGA codon may function as stop in *Naiad-Mmer*. Overall, the ORFs of each of the four families apparently encode selenoproteins that incorporate multiple Sec residues. Furthermore, following the phylogenetic relationship of *Naiads* presented in Fig. 5A, it is likely that the evolutionary transition to selenoproteins has occurred independently in spiders and clams.

### Structurally diverse Chlamys elements in the green lineage and protists

As part of a recent annotation of TEs in the unicellular green alga *Chlamydomonas reinhardtii* and its close relatives (Craig, et al. 2021), we identified novel PLE families with N-terminal GIY-YIG domains. These elements were termed *Chlamys*, although they were not further described. As with *Naiads*, the N-terminal EN+ PLEs in *Chlamydomonas* possess several of the defining features of canonical C-terminal EN+ PLEs, including genome-wide distributions, frequent 5’ truncation and partial tandem insertions producing pLTRs. As introduced in the following text, *Naiad* and *Chlamys* share several features and collectively form a strongly supported N-terminal EN+ clade (Fig. 5A, Fig. 8), although *Chlamys* elements also possess characteristics that distinguish them from the newly described metazoan clade.

The predicted proteins of the *Chlamys* elements included the *Naiad*-specific CxC zf-CDGSH-like Zn-finger motif and the IFD (Fig. S1, S4). Additionally, all but two *Chlamys* elements (*Chlamys-2_cBra* and *PLE-X1_cBra*) lacked HHRs, further strengthening their evolutionary link to *Naiads*. As before, we used these conserved features to perform an extensive search for related PLEs in other taxa. We curated *Chlamys* elements from a wide diversity of green algae, including species from the Chlorophyceaen order Sphaeropleales and the unicellular streptophyte algae *Mesostigma viride* and *Chlorokybus atmophyticus*. The Sphaeropleales and Chlamydomonadales are estimated to have diverged in the pre-Cambrian, while chlorophytes and streptophytes (which includes land plants) possibly diverged more than 1 billion years ago (Del Cortona et al., 2020). Additional curation identified more distantly related and structurally diverse families in the chlorophyte *Botryococcus braunii* (class Trebouxiophyceae), the multicellular streptophyte alga *Chara braunii*, two species of spike moss (genus *Selaginella*) and the myxomycete slime mold *Physarum polycephalum* (phylum Amoebozoa). Certain *Chlamys* families were also found in very high copy numbers, most notably in the genomes of the streptophytes *M. viride* and *C. braunii* (Fig. 5B).

*Chlamys* elements were mostly longer (3.3 – 8.2 kb, not including pLTRs) and more structurally diverse than *Naiads* (see below). The length of several families was also increased by the presence of tandem repeats. In the RT domain, the “DKG” motif was present as DK without a well-conserved third aa, and the IFD was generally longer (~20-40 aa) than that of *Naiads* (Fig. S1, S4). Targeted insertion at (CA)_n_ repeats was observed for many *Chlamys* elements. Relative to *Naiads*, the most striking difference was in the EN domain. Although the GIY-YIG EN is N-terminal in both *Chlamys* and *Naiads*, in *Chlamys* both the linker domain harboring the CX_2-5_CxxC Zn-finger and the CCHH Zn-finger motif are absent (Fig. S2). Thus, the EN of *Chlamys* differ from both *Naiads* and canonical C-terminal EN+ PLEs (all of which encode the CCHH motif, with the CX_2-5_CxxC Zn-finger absent in *Penelope/Poseidon*). The conserved R, H, E and N aa beyond the GIY-YIG core are all present in *Chlamys*. Finally, *Naiads* formed a well-supported clade to the exception of all *Chlamys* elements (Fig. 5A). Collectively, *Naiad* and *Chlamys* are distinguished based on both taxonomic and structural features, and they can be considered as two major ancient groups that together comprise a wider N-terminal EN+ clade.

The minimal *Chlamys* domain organization, which is shared by most families in *Chlamydomonas,* the Sphaeropleales and the unicellular streptophytes, is represented by *Chlamys-1_meVir* in Fig. 6. Five families from *Chlamydomonas* encoded proteins with plant homeodomain (PHD) finger insertions, which were either located between RT2a and RT3(A) (Fig. 7) or between the H and E conserved aa within EN. PHD fingers have been reported from TEs including *CR1* non-LTR elements (Kapitonov and Jurka 2003) and *Rehavkus* DNA transposons (Dupeyron, et al. 2019), where they may play a role in chromatin restructuring. PHD fingers are present in several other retrotransposons and DNA transposons in *C. reinhardtii*, and it appears to be a common accessory domain (Perez-Alegre, et al. 2005; Craig 2021). Several additional domains were encoded by the more distantly related *Chlamys* elements. Two divergent organizations were observed in *P. polycephalum* families, the first of which included an SAP domain inserted between RT7(E) and the RT thumb (*Chlamys-1_pPol*, Fig. 6). SAP (SAF A/B, Acinus and PIAS) is a putative DNA-binding domain that has previously been reported in *Zisupton* DNA transposons (Böhne, et al. 2012). The second type included the element *Physarum_Pp1,* which was first described from a 5’ truncated consensus as an unusual PLE with an IFD (Gladyshev and Arkhipova 2007). Extending the consensus sequence revealed a predicted protein with a reduced N-terminus that entirely lacked the CxC motif present in all other *Chlamys* and *Naiad*, and included a reduced EN domain in which the GIY-YIG motif was present but weakly conserved and the region containing the conserved R, H, E and N aa was absent (Fig. 3A, 6). Although the *P. polycephalum* genome is highly fragmented, *Physarum_Pp1* does appear to be present genome-wide.

**Fig. 6.**
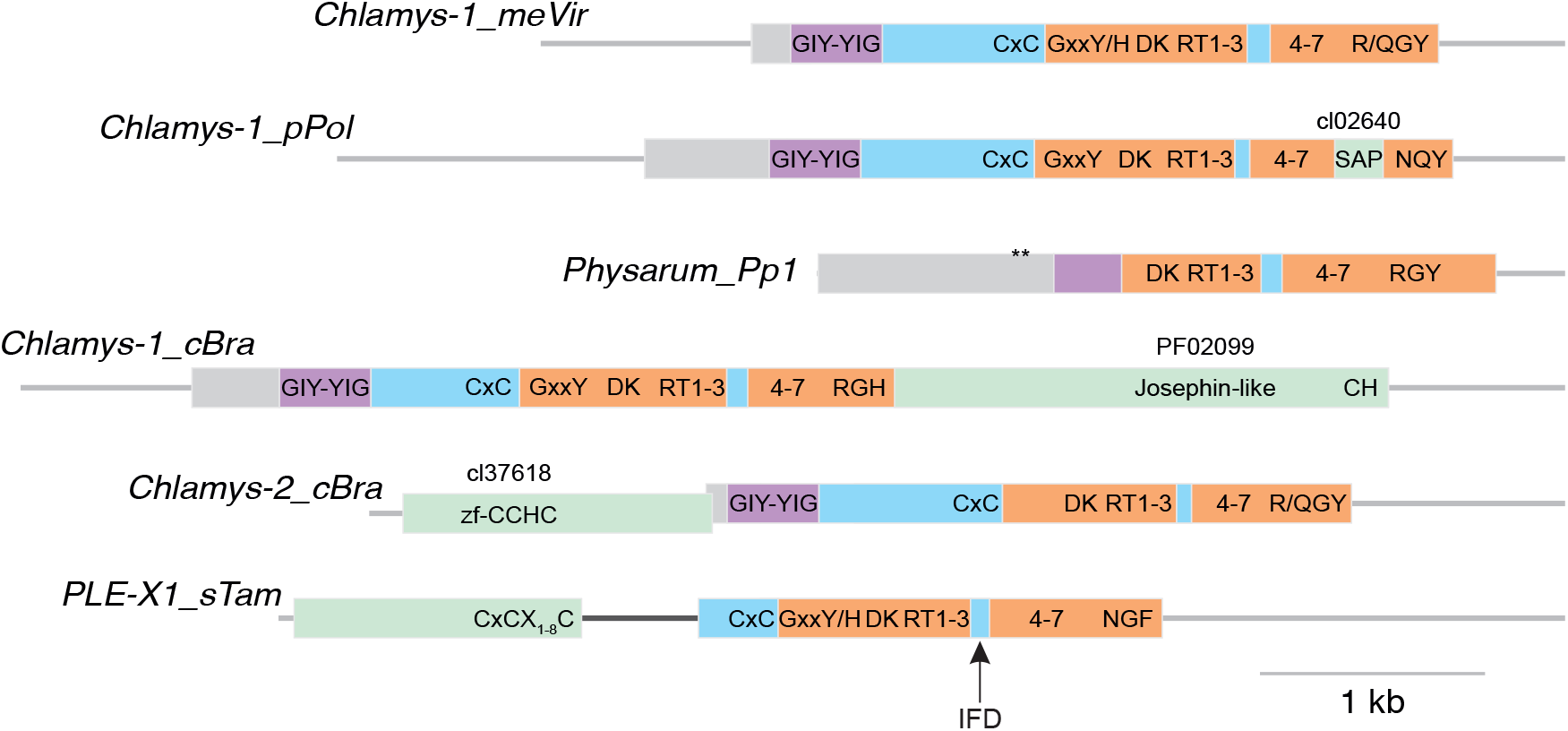
Structural diversity of *Chlamys* elements. Domain architecture of *Chlamys* elements is represented by to scale schematics. Thin gray lines represent sequences not present in ORFs. Domains are colored as follows: RT, peach; GIY-YIG EN, purple; insertions/extensions containing conserved motifs, green; N-terminal extensions without recognized domains, gray; regions specific to *Naiad* and *Chlamys*, light blue. Purple is used for the GIY-YIG EN to distinguish the *Chlamys* EN from the *Naiad* EN, which contains the CCHH motif and is represented in yellow in Fig. 1. The most conserved amino acid motifs and the highest-scoring PFAM/CD domain matches are also shown. The asterisks on the *Physarum_Pp1* model represent in-frame stop codons, which may indicate the presence of an undetected intron. Note the *Physarum_Pp1* EN-like domain is also reduced and weakly conserved (Fig. 3A). The dark gray line in *PLE-X1_sTam* represents an intron that was inferred from *S. moellendorffii* annotated gene models.

**Fig. 7.**
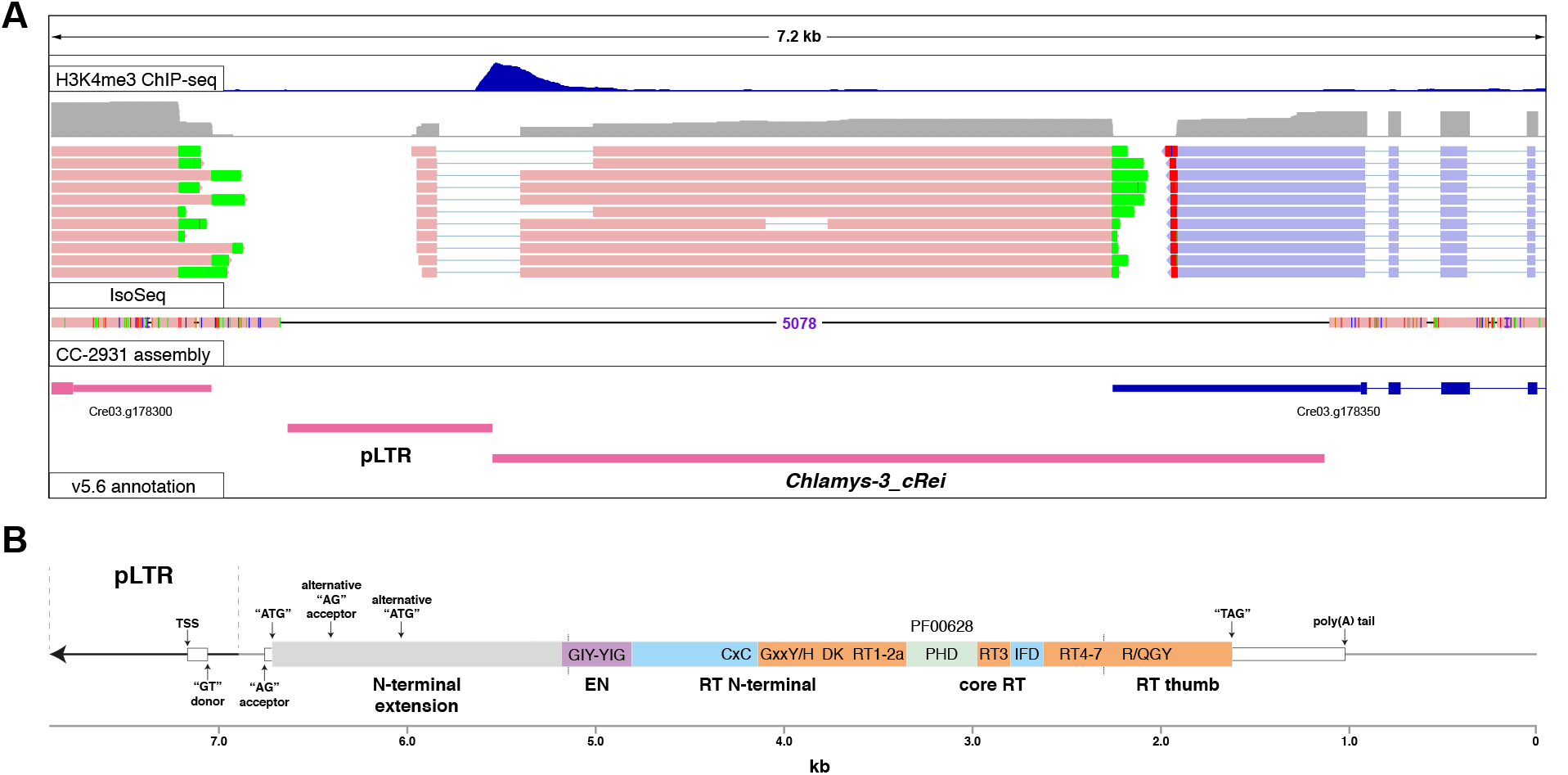
Functional characterization of an active *Chlamys* element in *C. reinhardtii.* **(A)** IGV browser view (Robinson, et al. 2011) of a *Chlamys-3_cRei* copy that is polymorphic between the reference genome and the divergent field isolate CC-2931. Green and red mismatched bases on Iso-Seq reads represent poly(A) tails of transcripts. Forward strand reads and gene/repeat models are shown in pink, and reverse strand in blue. **(B)** Schematic of the inferred gene model and structural organization of *Chlamys-3_cRei*. Note that this represents the full-length element and the transcribed copy above contains a 2.84 kb internal deletion, the boundaries of which are shown by the dashed black lines. Domains are colored as in Fig. 6 and conserved motifs are textually represented as shown for *Chlamys-1_meVir* in that figure.

EN+ PLEs were reported from the *C. braunii* genome project, although Nishiyama, et al. (2018) did not further describe these elements. We observed three distinct types of *Chlamys* in *C. braunii*. The first possessed long ORFs (~1,800 aa) encoding peptides with a C-terminal extension including a motif with weak homology to Josephin and a second motif with several well-conserved C and H aa (*Chlamys-1_cBra*, Fig. 6). Josephin-like cysteine protease domains are present in *Dualen* non-LTR elements, where they may play a role in disrupting protein degradation (Kojima and Fujiwara 2005). The second type included an upstream ORF encoding a peptide with a gag-like zinc-knuckle domain (zf-CCHC, *Chlamys-2_cBra*, Fig. 6). The third type was notable since related elements were also identified in spike mosses, and a small number of highly significant BLASTp results were recovered from moss species, potentially indicating a wider distribution in “early-diverging” plants. These families include the CxC motif but lack the GIY-YIG EN, with a unique N-terminal extension that is likely separated by an intron and includes a conserved CxCX_1-8_C motif (*PLE-X1_sTam*, Fig. 6). The families in *C. braunii* appeared to have genome-wide distributions, and remarkably, the two spike moss families exhibited targeted insertions at a precise location within 28S ribosomal RNA genes. The insertion target differed by only 4 bp between the families (Fig. S5), suggesting deep conservation of the target sequence at least since the divergence of *S. moellendorffii* and *S. tamariscina* ~300 Mya (Xu, et al. 2018). Metazoan ribosomal DNA is a well-documented insertion niche for *R* element non-LTRs and the *piggyBac* DNA transposon *Pokey* (Eickbush and Eickbush 2007), although to our knowledge this is the first example from both plants and PLEs. It remains to be seen how this group achieve either genome-wide or targeted ribosomal DNA insertion without an identified EN. Interestingly, these families form a well-supported clade with the EN+ family from *P. polycephalum* (*Chlamys-1_pPol*, Fig. 5A), potentially indicating secondary loss of the GIY-YIG EN.

### Functional characterization of an active Chlamys element

As high-quality functional data is available for *C. reinhardtii*, we further focused on the 10 *Chlamys* families curated in this species. Notably, we also identified putatively nonautonomous *Chlamys* elements, which produced pLTRs (and often multi-copy head-to-tail insertions) and generally exhibited sequence similarity to autonomous families at their 3’ ends. The nonautonomous elements include *MRC1*, which was previously described as a nonautonomous LTR (Kim, et al. 2006) and may be the most active TE in *C. reinhardtii* laboratory strains (Neupert, et al. 2020). Further supporting recent activity, *Chlamys* copies exhibited minimal divergence from their respective consensus sequences (Fig. S6) and within-species polymorphic insertions were observed for copies of all 10 autonomous families by comparison to a newly assembled PacBio-based genome of the divergent field isolate CC-2931 (Supp. note). Cumulatively, *Chlamys* PLEs spanned ~1.6% of the 111 Mb *C. reinhardtii* genome and comprised ~15% of the total TE sequence.

Only one active C-terminal EN+ PLE has been experimentally characterized, the archetypal *Penelope* of *D. virilis* (Pyatkov, et al. 2004; Schostak, et al. 2008). In an attempt to characterize a *Chlamys* element, we searched for an actively transcribed copy using recent PacBio RNA-seq (i.e. Iso-Seq) and H3K4me3 ChIP-seq datasets (Gallaher, et al. 2021), with the H3K4me3 modification reliably marking active promoters in *C. reinhardtii* (Ngan, et al. 2015). Due to frequent 5’ truncation, only two families were found with full-length copies, and transcription was observed for only a single copy of the *Chlamys-3_cRei* family (Fig. 7A). Unfortunately, this copy features a 2.8 kb deletion, although this is entirely within the ORF and the copy presumably retains a functional promoter, transcription start site (TSS) and terminator. Strikingly, the derived gene model of *Chlamys-3_cRei* (Fig. 7B) shared several features with *Penelope*, in which the pLTR harbors the TSS and a 75 bp intron within the 5’ UTR that overlaps the internal promoter (Arkhipova, et al. 2003; Schostak, et al. 2008). In *Chlamys-3_cRei*, the TSS is also located in the pLTR and a 398 bp intron within the 5’ UTR spans the boundary between the pLTR and downstream main body. The H3K4me3 ChIP-seq supports an internal promoter coinciding with the intron. Additionally, three Iso-Seq reads supported an alternative isoform with a 751 bp intron. This isoform initiates at a downstream start codon and results in a peptide truncated by 293 aa, although as the predicted *Chlamys-3_cRei* peptide includes an N-terminal extension both isoforms encode complete EN and RT domains. The similarities between *Penelope* and *Chlamys-3_cRei* potentially indicate an ancient and deeply conserved organization and perhaps mechanism shared by canonical PLEs and the N-terminal EN+ PLEs described herein.

### Hydra: A novel C-terminal EN+ clade

While performing an updated phylogenetic analysis of all PLEs (see below), we noticed that seven C-terminal EN+ families in Repbase formed an isolated group highly divergent from *Neptune, Penelope/Poseidon* and *Nematis*. All but one of these families were annotated from the freshwater polyp *Hydra magnipapillata*. Using protein homology searches, we identified a small number of additional families in other aquatic invertebrates spanning four phyla (Cnidaria, Mollusca, Echinodermata, and Arthropoda), notably in species such as the stony coral *Acropora millepora* and the sea cucumber *Apostichopus japonicus*. These elements were generally short (<3 kb) and contained single ORFs encoding peptides with several similarities to canonical C-terminal EN+ PLEs, i.e. no CxC motif, no IFD and a C-terminal GIY-YIG EN (Fig. S7). HHRs were also detected, strengthening the relationship with canonical PLEs. However, these families also exhibited unique features. The N-terminal GxKF/Y motif was not well conserved, the DKG motif was modified to DKT, and RT4(B) was particularly divergent and challenging to align. Most notably, in the EN domain the CCHH motif universal to C-terminal EN+ PLEs (and *Naiads*) was absent (Fig. S2, S7). A linker domain was present which was most similar to that of *Nematis*, although the CX_2-5_CxxC Zn-finger was modified to a CxCX_5_C motif. Interestingly, all families exhibited insertions into (TA)n, strengthening the association between the linker domain and targeted insertion. We name this new clade of C-terminal EN+ PLEs *Hydra*, in line with both their aquatic hosts and their discovery in *H. magnipapillata*.

### Evolution of the RT and GIY-YIG EN domains

As seen in Fig. 5, the newly discovered types of PLE span much of the well-sequenced taxonomic diversity in Eukarya, including protists, plants, and animals. We placed *Naiad, Chlamys* and *Hydra* representatives into a reference PLE dataset that included the previously known EN+ *Penelope/Poseidon, Neptune, Nematis* and EN–*Athena* and *Coprina* clades, as well as representatives of the sister clade to PLEs, the TERTs (Arkhipova 2006; Gladyshev and Arkhipova 2007). The combined phylogeny of the extended core RT domain, which also includes the RT thumb and the previously identified N-terminal conserved motifs N1-N3 (Arkhipova 2006), is presented in Fig. 8. With the exception of *Neptune*, all of the above clades were recovered with ultrafast bootstrap support values >90%. The monophyly of *Neptune* was not recovered, with *Neptune* elements forming a paraphyletic group in a weakly supported clade with the taxonomically diverse EN–*Coprina* elements, as also occurred in a previous analysis (Arkhipova, et al. 2017). The novel N-terminal EN+ elements (i.e. *Naiad* + *Chlamys*) formed a strongly supported clade, although *Chlamys* was paraphyletic with respect to the *Naiad* clade and the internal topologies of the more structurally diverse *Chlamys* elements were not well supported. Despite its potential paraphyly, we still consider *Chlamys* to be a useful grouping given its unique structural features. The rotifer-specific EN–*Athena* elements formed the most basal PLE clade when rooting the phylogeny on TERTs, although the deep branches linking the major PLE clades generally received weak support. As seen in both *Chlamys* and *Neptune*, EN–families occasionally emerge within EN+ clades, apparently as a result of EN loss accompanied by acquisition of an alternative way of employing accessible 3’-OH ends for RT priming. Overall, it is evident that, with inclusion of the hitherto unknown superfamilies, PLE RTs display an astonishing level of clade diversity, which is comparable to that of non-LTR and LTR retrotransposon RTs, and will undoubtedly increase in line with the number of sequenced genomes from under-represented eukaryotic branches of the tree of life.

**Fig. 8.**
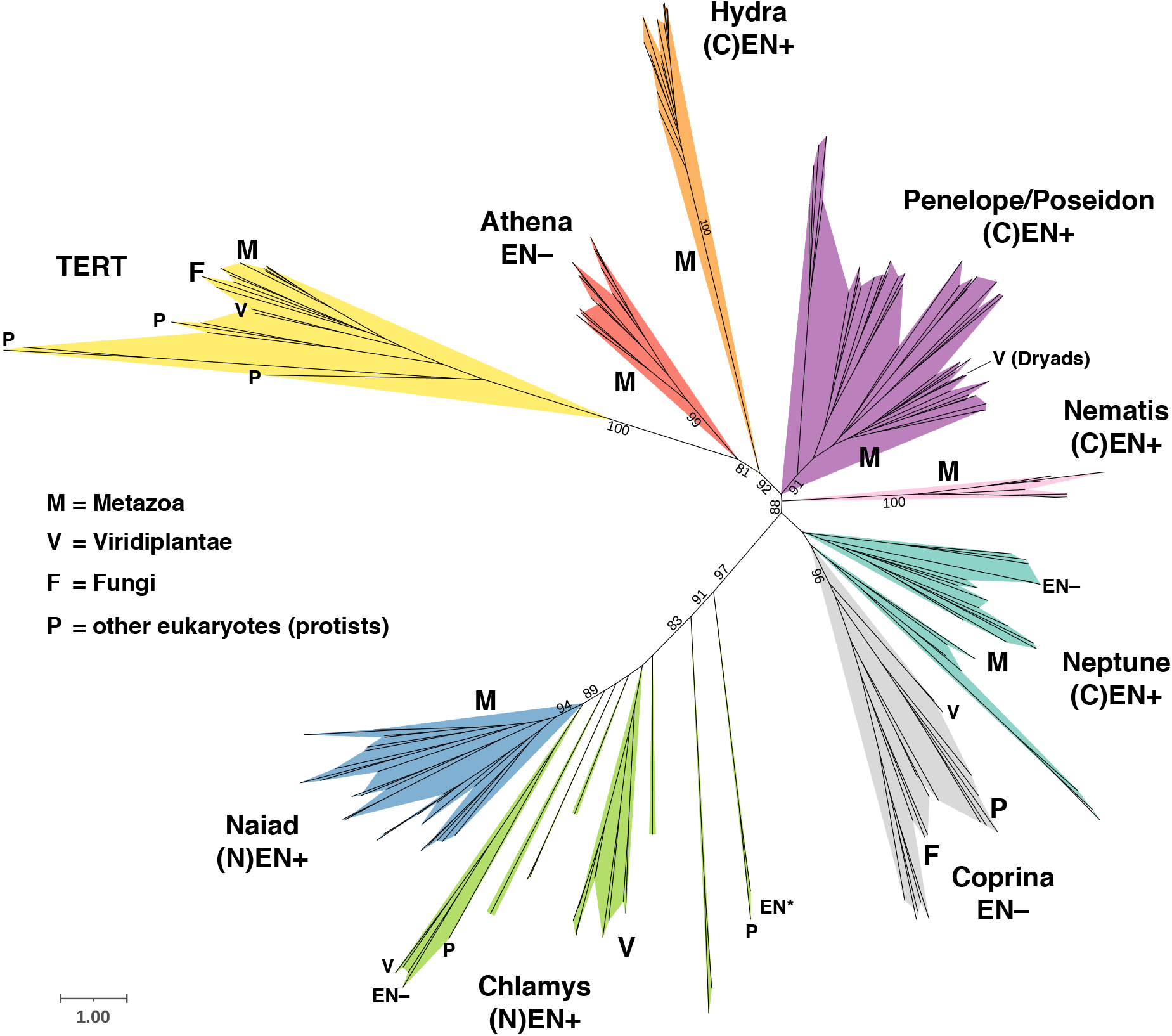
Core RT maximum likelihood phylogeny of PLE RTs and TERTs. Support values from 1000 ultrafast bootstrap replications are shown, with all values from nodes within major clades (colored) and any values <70 at deeper nodes excluded to aid visualization. The taxonomic range of clades and subclades is shown by letters. Note that the “V” marking the “*Dryad*” subclade within *Penelope/Poseidon* points at three conifer families from the presumed horizontal transfer event (Lin, et al. 2016). The location of EN in EN+ groups is provided by the prefixes (N) and (C) for N-terminal and C-terminal, respectively. Subclades with EN remnants or no EN that are within EN+ clades are shown by EN* and EN–tags, respectively. Scale bar, aa substitutions per site. The phylogeny was annotated using iTOL (Letunic and Bork 2019).

In light of the increased diversity uncovered by *Naiad, Chlamys* and *Hydra* PLEs, we also attempted to further elucidate the evolutionary relationships of the GIY-YIG EN domain. Since the domain is too short for conventional phylogenetic analysis, it has previously been analyzed using protein clustering approaches. Dunin-Horkawicz, et al. (2006) found that the most similar ENs to those from canonical C-terminal EN+ PLEs belonged to the HE_Tlr8p_PBC-V_like group (cd10443), which includes homing ENs from bacteria, chloroviruses (e.g. *Paramecium bursaria* chlorella virus 1, PBCV-1) and iridoviruses, as well as an EN from the *Tlr8* Maverick/Polinton element from *Tetrahymena thermophila*. Using CLANS (Frickey and Lupas 2004), we performed an updated clustering analysis with all available PLE ENs (Fig. S8). *Neptune, Nematis* and *Penelope/Poseidon* ENs formed distinct although strongly connected clusters, with *Naiad* ENs essentially indistinguishable from *Neptune*. These results largely follow expectations from the shared presence of the CCHH motif and the presence/absence of the CX_2-5_CxxC Zn-finger linker (Fig. S2), and these domains are collectively representative of the canonical PLE EN described in NCBI (cd10442). *Hydra* ENs formed a distinct and well-resolved cluster that was nonetheless related to other PLE ENs, in line with the absence of the CCHH motif and the alternative configuration of the linker motif. Interestingly, the ENs sometimes associated with the giant *Terminons* (Fig. 1), which contain RTs from the otherwise EN–*Athena* group (Arkhipova, et al. 2017), were also recovered as a distinct cluster related to other PLE ENs. These ENs include both the CX_2-5_CxxC and CCHH motifs, suggesting shared ancestry with EN+ PLEs, although they also contain large unique insertions (Fig. S2). Finally, the *Chlamys* ENs were diffusely clustered between all other PLE ENs and several ENs from the HE_Tlr8p_PBC-V_like group. The lack of strong clustering can likely be explained by the lack of both the CX_2-5_CxxC and CCHH motifs resulting in fewer conserved sites, and the possible link between *Chlamys* and HE_Tlr8p_PBC-V_like ENs should be interpreted tentatively. Overall, the GIY-YIG ENs of all PLEs appear to be related, and in line with the results of Dunin-Horkawicz, et al. (2006), PLE ENs are most similar to particular homing ENs from bacteria and viruses.

## DISCUSSION

### A new major PLE clade with N-terminal EN and its impact on genome and transposon annotation

*Penelope*-like elements are arguably the most enigmatic type of retrotransposable elements inhabiting eukaryotic genomes. Due to their absence from the best-studied genomes such as mammals, birds and angiosperms, and the complex tandem/inverted structures brought about by still undefined features of their peculiar transposition cycle, PLEs have largely been neglected and overlooked by most computational pipelines used in comparative genomics. Current approaches distinguish PLEs by the presence of a PLE-related RT, and classify them to the “order” level as a clade of non-LTR elements (Bao, et al. 2015) without subdivision into groups differing by domain architecture and phylogenetic placement, as is commonly done for non-LTR (LINE) and LTR retrotransposons (Storer, et al. 2021). Here we show that the degree of PLE structural and phylogenetic diversity matches that of non-LTR and LTR retrotransposons, emphasizing the need for updating current classification schemes and TE-processing computational pipelines.

Our data also underscore the need to adjust computational pipelines to incorporate searches for GIY-YIG EN either upstream or downstream from PLE RT, due to the high degree of polymorphisms (especially frameshifts) in the connector region, which complicates identification of full-length elements. This is especially relevant at a time when increasing numbers of invertebrate genomes are being sequenced, with *Naiad* elements often contributing tens of thousands of copies to metazoan genomic DNA. Under-annotation of poorly recognizable TEs poses a serious problem to gene annotation. This is especially well-illustrated in host-associated and environmental metagenome analyses, where understudied eukaryotic TEs become mis-assigned to bacterial genomes and are propagated in taxonomy-aware reference databases, jeopardizing future automated annotations (Arkhipova 2020).

Of special interest is the dominance of *Chlamys* PLEs in the plant kingdom, where their ancient nature is supported by their presence in the most basal members of the green lineage, by a high degree of divergence between *Chlamys* elements, and by distinctive features of the associated EN. In contrast to the documented case of horizontal transfer of a canonical C-terminal EN+ PLE into conifer genomes (Lin, et al. 2016), their early-branching position in the PLE phylogeny argues that they constitute ancestral genome components in early-branching plants and green algae, and does not support recent introduction. Nevertheless, their ongoing activity and diversification in *Chlamydomonas* indicates that *Chlamys* elements are actively participating in algal genome evolution.

### *Common and distinctive features of Naiad* and *Chlamys retrotransposition*

Consistent association of all PLE RTs with a special type of endonuclease/nickase (GIY-YIG EN), which may have occurred several times in early eukaryotic evolution to form distinct lineages characterized by N- or C-terminal EN domains, underscores the importance of this EN for efficient intragenomic proliferation mediated by PLE RT, and emphasizes the need for further mechanistic investigations of the non-trivial PLE transposition cycle in representatives of each PLE lineage. It is very likely that the unique EN cleavage properties determine the formation of complex tandem/inverted pLTRs and the “tail” extension on either side of PLEs, not observed during TPRT of non-LTR elements, but seen in *Naiad/Chlamys*.

Further, PLEs are highly unusual among retroelements in their ability to retain introns after retrotransposition, sometimes even retrotransposing intron-containing host genes *in trans* (Arkhipova, et al. 2003; Arkhipova, et al. 2013). While most of the *Naiad/Chlamys* ORFs are not interrupted by introns, the functionally characterized active *Chlamys-3_cRei* element shares an intron position within the 5’-UTR with the functionally studied *Penelope* from *D. virilis*, overlapping with the internal promoter (Schostak, et al. 2008). This suggests that other PLEs may share this organization and harbor introns upstream of the main ORF. The significance of intron retention is unknown, although it is likely a consequence of the unusual retrotransposition mechanism.

Interestingly, we did not detect HHR motifs in *Naiad* or *Chlamys* elements, except for *Chlamys-2_cBra* (EN+) and *PLE-X1_cBra* (EN–). These two families from *C. braunii* are not closely related (Fig. 5A), implying HHRs may have been independently acquired or frequently lost from other *Chlamys*. Conversely, HHRs are universally present in the *Hydra* clade and other PLEs. HHR function in EN+ PLEs is still unclear, and while they have been hypothesized to help cleave the tandemly arranged long precursor RNAs (Cervera and De la Peña 2014), their absence from *Naiads* and most *Chlamys* elements obviously does not interfere with their successful intragenomic proliferation.

In many cases, it was not possible to discern target-site duplications in *Naiads* and *Chlamys* due to a strong insertion bias towards microsatellite repeats, with (CA)_n_ most commonly observed in *Chlamys* and (TA)_n_ in *Naiads*. The CX_2-5_CxxC EN linker was hypothesized to mediate such bias in *Neptune* PLEs (Arkhipova 2006) and could do so in *Naiads*, but its absence from *Chlamys* suggests that the novel CxC domain may also play a role in targeting EN activity to specific DNA repeats. Also of interest are the EN– “*PLE-X*” families from two species of spike moss, which are the first known TEs to exhibit targeted insertion into the 28S ribosomal RNA gene in plants, as is observed in certain non-LTRs and DNA transposons of arthropods and other animals (Eickbush 2002; Penton and Crease 2004; Gladyshev and Arkhipova 2009).

Finally, it is unknown what role the IFD may play in *Naiads* and *Chlamys*. In TERTs, the IFD aids the stabilization of telomerase RNA (TER) and DNA during the extension of telomeric DNA (Jiang, et al. 2018). The IFD domain in *Naiads* and *Chlamys* is shorter than that of TERTs, and its loss from a specific *Naiad* subclade demonstrates that it is not necessarily a functional requirement.

Establishment of an *in vitro* system to study PLE retrotransposition mechanisms would be the next important task required to achieve full understanding of PLE-specific TPRT features that distinguish them from LINEs, such as formation of complex tandem/inverted repeat structures and microsatellite insertion bias.

### Naiad selenoproteins

The *Naiads* that encode selenoproteins are notable for two reasons. First, almost all described selenoproteins include a single Sec, whereas the *Naiads* contain either three or four. Baclaocos, et al. (2019) performed analysis of selenoprotein P (SelP), one of the few selenoproteins including multiple Sec residues, finding that in bivalves SelP contains the most Sec residues of any metazoan group, and that spider SelP proteins contain a moderate number of Sec residues. Bivalves in particular are known for their high selenium content (Bryszewska and Måge 2015), and it may be that the *Naiads* represent cases of TEs adapting to their host cellular environments. However, even in bivalves selenoproteins are incredibly rare (e.g. the pacific oyster selenoproteome encompasses 32 genes (Baclaocos, et al. 2019)), suggesting a more specific role for the incorporation of Sec in these families. Sec residues are involved in numerous physiological processes and are generally found at catalytic sites, where in many cases they have a catalytic advantage relative to cysteine (Labunskyy, et al. 2014). All but one of the Sec residues in *Naiad* peptides correspond to highly conserved sites in the CX_2-5_CxxC Zn-finger, CCHH Zn-finger and the DKA motif, and although the precise physiological role of these motifs in PLEs is unknown, it may be that the incorporation of Sec provides both a catalytic and evolutionary advantage.

Second, the *Naiad* families are the first described selenoprotein-encoding TEs. It is currently unclear whether these represent highly unusual cases, although the fact that they appear to have evolved independently in spiders and clams hints that other examples may be found in the future. This has potential implications for TE annotation in general, and selenoprotein-encoding TEs may have previously been overlooked in taxa such as bivalves because of apparent stop codons. Additionally, this result may provide insight into evolution of new selenoproteins. The transition from encoding Cys to Sec is expected to be a complex evolutionary process, since a gene must acquire a

SECIS element and near-simultaneously undergo a mutation from TGT/TGC (encoding Cys) to TGA (Castellano, et al. 2004). The insertion of TEs carrying SECIS elements into the 3’ UTRs of genes could provide a pathway for SECIS acquisition, especially for TEs that undergo 5’ truncation and may insert with little additional sequence. It remains to be seen if the selenoprotein-encoding *Naiads*, or indeed any other TEs, have contributed to the evolution of new selenoproteins in their host genomes.

### Evolutionary implications for PLE origin and diversification

As the branching order of major PLE clades diverging from TERTs is not exceptionally robust, it may be difficult to reconstitute evolutionary scenarios which were playing out during early eukaryogenesis. It is possible that an ancestral EN–PLE, similar to *Athena* or *Coprina* but lacking the extended N-terminus, was present at telomeres (Gladyshev and Arkhipova 2007) before undergoing either multiple domain fusions to give rise to TERTs, or fusions with GIY-YIG EN, either at the N- or at the C-termini, to form the contemporary *Naiad/Chlamys, Neptune, Nematis*, and *Penelope/Poseidon* superfamilies capable of intrachromosomal proliferation.

There are several plausible evolutionary scenarios that could explain the observed EN and RT diversity, and ENs may have been acquired or exchanged several times by different PLE clades. It is possible that *Chlamys* elements acquired an EN without the CX_2-5_CxxC and CCHH motifs from a homing EN from the HE_Tlr8p_PBC-V_like family, and that the Zn-finger motifs were later gained by *Naiads*. ENs with both Zn-fingers could then have been transferred from *Naiads* to the C-termini of EN–animal PLEs (once or multiple times), giving rise to other EN+ clades. This scenario would imply that the internal CCHH was then lost in *Hydra*, and the upstream linker domain was either reduced (*Nematis*), reduced and modified (*Hydra*) or lost (*Penelope/Poseidon*). Alternatively, an EN containing one or both Zn-fingers could have been independently acquired by C-terminal EN+ PLEs (again once or multiple times) and exchanged with *Naiads* replacing the *Chlamys-*like EN (or gained independently by *Naiads* from a similar homing EN). This scenario would imply the existence of homing ENs with Zn-finger motifs, which have not been found, however both the CX_2-5_CxxC and CCHH motifs are present in the stand-alone ENs occasionally associated with *Terminons*. EN acquisition, either at the N- or C-terminus, may have been facilitated if RT and EN were brought in proximity either on a carrier virus or on a chimeric circular replicon allowing permutation. Any combination of events in the above scenarios could of course explain the observed diversity. Notably, early metazoans such as cnidarians exhibit the highest PLE clade diversity, with *Poseidon, Naiad* and *Hydra* present in *H. magnipapillata* and *Neptune, Naiad* and *Hydra* in the coral *A. millepora*, implying that the appropriate conditions existed for either multiple exchanges or acquisitions of ENs. Finally, EN losses are not unusual, and EN–elements can emerge within EN+ clades, as in *Chlamys* PLE-X families in *Selaginella* and *Chara*, or the *Neptune*-like *MjPLE01* from the kuruma shrimp *Marsupenaeus japonicus* (Koyama, et al. 2013), if they adopt alternative means of securing 3’-OH groups for TPRT.

While the IFD domain may have been inherited by *Naiads/Chlamys* from a common ancestor with TERTs, IFD-like regions are also found sporadically in *Coprina* elements, arguing against its use as a synapomorphy. It is possible that the IFD has been lost multiple times in different PLE clades, as demonstrated by its loss in some *Naiads*, or gained independently in *Naiads/Chlamys* and *Coprina*. The presence/absence of HHR motifs does not provide many clues either: while found in only two basal *Chlamys*-like elements and absent from *Naiads*, they are present in all other PLEs, both EN+ and EN–. As with newly described retrozymes, they may exploit autonomous PLEs for their proliferation (Cervera and de la Peña 2020), or they could provide an unknown function.

Regardless of the exact sequence of events which led to PLE diversification in early eukaryotic evolution, it is now clear that the diversity of PLE structural organization, manifested in the existence of at least seven deep-branching clades (superfamilies) differing by domain architecture and found in genomes of protists, fungi, green and red algae, plants, and metazoans from nearly every major invertebrate and vertebrate phylum, can no longer be overlooked and should be reflected in modern genomic analysis tools. As more genomes from neglected and phylogenetically diverse lineages become available, it is likely that the diversity of PLEs will continue to expand, further supporting their increasingly important and unique position in TE biology and their contribution to shaping the amazing diversity of eukaryotic genomes.

## MATERIALS AND METHODS

### Annotation and curation of PLE consensus sequences

For general TE identification and annotation in metazoan genome assemblies (*D. stevensoni, P. laevis*), we used TEdenovo from the REPET package (Flutre, et al. 2011) to build *de novo* repeat libraries with default parameters. Although REPET-derived *de novo* TE consensus sequences are automatically classified under Wicker’s scheme (Wicker, et al. 2007), we additionally used RepeatMasker v4.1.0 (Smit, et al. 2015) for TE classification, detection and divergence plot building, using the initial TEdenovo repeat library. To specifically illustrate the composition on PLE families in the *P. laevis* genome, we used the corresponding consensus sequences of PLE families as a local library for divergence plot building.

Initial *Chlamys* consensus sequences from *C. reinhardtii* and its close relatives (*Chlamydomonas incerta*, *Chlamydomonas schloesseri* and *Edaphochlamys debaryana*) were curated as part of a wider annotation of TEs in these species (Craig 2021; Craig, et al. 2021). Inferred protein sequences from the metazoan and algal consensus models were then used as PSI-BLAST or tBLASTn (Camacho, et al. 2009) queries to identify related *Naiad* and *Chlamys* PLEs in other species. PSI-BLAST was run using NCBI servers to identify putative PLE proteins that had been deposited in NCBI. tBLASTn was performed against all eukaryotic genome assemblies accessed from NCBI on 2020/04/09. Assemblies with multiple significant hits were selected for further curation, and where several related species had multiple hits the most contiguous assemblies were targeted. A Perl script was used to collect the nucleotide sequence of each tBLASTn hit from a given assembly, and the most abundant putative PLEs in each species were subjected to manual curation. This was performed by retrieving multiple copies by BLASTn, extending the flanks of each copy and aligning the subsequent sequences with MAFFT v7.273 (Katoh and Standley 2013). The multiple sequence alignment of each family was then visualized and manually curated (removing poorly aligned copies, identifying 3’ termini and pLTRs if present, etc.). Consensus sequences were produced for each family and protein sequences were inferred by identifying the longest ORF.

Copy number was estimated by performing BLASTn against the assembly using the consensus sequence as a query. NCBI BLASTn optimized for highly similar sequences (megablast) was used with cutoff E-value 1e-5. Whole-genome shotgun (wgs) datasets were used for each species, with the best quality assembly used in case of multiple isolates. In most cases, NCBI web interface was used to control for truncated and deleted copies via graphical summary. If the maximum number of target sequences (5000) was exceeded, wgs datasets were created using blastn_vdb from the SRA Toolkit and searched with blastn 2.6.1+, or installed locally and searched with blastn 2.10.1+.

Novel *Hydra* families were identified and curated using the same approach as above. Existing protein sequences from *H. magnipapillata* PLEs accessed from Repbase were used as initial queries to search for related elements, alignments of which were then manually curated and used to produce consensus sequences.

### Functional motif identification

SECIS elements were identified in *Naiad* consensus sequences containing in-frame UGA codons using the SECISearch3 (Mariotti, et al. 2013) online server (http://gladyshevlab.org/SelenoproteinPredictionServer/).

Hammerhead ribozyme motif (HHR) motif searches were performed using secondary structure-based software RNAmotif (Macke, et al. 2001). A general HHR descriptor (Cervera and De la Peña 2014) was used to detect HHR motifs in *Naiad/Chlamys* and *Hydra* elements. More relaxed descriptors were also employed as in Arkhipova, et al. (2017) to accommodate different helices with longer loops and stem mispairing and more relaxed cores with mismatches, and with and without the presence of Helix III, however it did not result in additional HHR motif detection.

### Functional characterization of Chlamys elements in Chlamydomonas reinhardtii

The divergence landscape (Fig. S6) and total abundance of *Chlamys* elements in *C. reinhardtii* were calculated using RepeatMasker v4.0.9 (Smit, et al. 2015) and the highly-contiguous assembly of strain CC-1690 (O’Donnell, et al. 2020). The functional characterization represented in Fig. 7 was performed using the standard v5 reference genome. Iso-Seq (accession: PRJNA670202) and H3K4me3 ChIP-seq (accession: PRJNA681680) data were obtained from Gallaher, et al. (2021). CCS (circular consensus sequence) Iso-Seq reads were mapped using minimap2 (-ax splice:hq – secondary no) (Li 2018). Within-species polymorphism was demonstrated by comparison to a *de novo* PacBio-based assembly of the divergent field isolate CC-2931, which exhibits ~3% genetic diversity relative to the standard reference strain (Craig, et al. 2019). The sequencing and assembly of the CC-2931 assembly is described in Supp. note. The CC-2931 assembly was mapped to the v5 reference using minimap2 (-ax asm10).

### RT phylogeny and EN protein clustering analysis

Initial amino acid sequence alignments were done with MUSCLE (Edgar 2004), with secondary structure assessed by inclusion of TERT PDB files (3kyl, 3du5) using PROMALS3D (Pei, et al. 2008), and were manually adjusted to ensure the presence of each conserved motif at the proper position. Phylogenetic analysis was done with IQ-TREE v1.6.11 (Trifinopoulos, et al. 2016), with the best-fit model chosen by ModelFinder according to Bayesian information criterion, and with 1000 UFBoot replicates to evaluate branch support.

Protein clustering of the GIY-YIG EN domain was performed using CLANS (Frickey and Lupas 2004). GIY-YIG ENs from all superfamilies annotated at NCBI (cd00719) were combined with those from PLEs (canonical C-terminal EN+, N-terminal EN+, *Hydra* and *Terminons*). All ENs were reduced to the core domain spanning from the “GIY” motif to the conserved N aa, unless a Zn-finger linker domain was present upstream of the “GIY”, in which case this motif was also included (see Fig. S2). Several very distantly related EN superfamilies were excluded after a preliminary analysis. CLANS was run with a p-value threshold of 1×10^−8^ until no further changes were observed in clustering.

## Acknowledgments

This work was supported by the U.S. National Institutes of Health grant R01GM111917 to I.A. R.C. was supported by the Biotechnology and Biological Sciences Research Council EASTBIO Doctoral Training Partnership and the project received funding from the European Research Council under the European Union’s Horizon 2020 Research and Innovation Programme (Grant Agreement no. 694212).

