## Supplemental Note for "An ancient clade of *Penelope*-like retroelements with permuted domains is present in the green lineage and protists, and dominates many invertebrate genomes"

### Supplemental Note: Sequencing and assembly methods for *Chlamydomonas reinhardtii* strain CC-2931

CC-2931 is a field isolate of *Chlamydomonas reinhardtii* sampled in North Carolina in 1991. The strain exhibits ~3% genetic diversity relative to the reference genome strain, and is likely part of a population that is at least partially reproductively isolated from that of the reference strain (Craig, et al. 2019). High molecular weight DNA was extracted from a 4d culture of CC-2931 following the protocol of Craig, et al. (2021). Genomic DNA was sequenced on the Pacific Biosciences (PacBio) Sequel platform at Edinburgh Genomics, yielding 7.51 Gb of reads with an N50 of 20.96 kb. A *de novo* contig-level assembly was produced using wtdbg2 (Ruan and Li 2020) using the parameters "-g 111m -x sq". PacBio-based polishing was performed with two iterations of Arrow (https://github.com/PacificBiosciences/GenomicConsensus), mapping reads with pbmm2 (https://github.com/PacificBiosciences/pbmm2). Illumina-based polishing was performed with one iteration of Pilon (Walker, et al. 2014) with the flag "--fix bases". Illumina data were obtained by the whole-genome re-sequencing of 14 mutation accumulation lines derived from CC-2931 (Ness, et al. 2015). These experimental lines were clonally maintained and allowed to accumulate mutations in order to study mutation rates and properties. Data from each line were subsampled to 10% coverage to ensure that no mutations, which are expected to be unique to single lines, were incorporated into the assembly. The assembly spanned 108.95 Mb on 177 contigs with an N50 of 3.01 Mb.

The polished contigs were then manually scaffolded to chromosomes by alignment to the CC-1690 assembly (O'Donnell, et al. 2020) using MashMap v2.0 (Jain, et al. 2018). Gaps between contigs were filled with 10 kb of "N" unknown bases. All regions where a CC-2931 contig mapped across two CC-1690 chromosomes were manually inspected against the raw PacBio reads using IGV v2.7.2 (Robinson, et al. 2011). All such breaks in synteny were strongly supported, which implied the presence of one large inversion and three reciprocal translocations in CC-2931 relative to laboratory strains (see Craig (2021)). Given the putative rearrangements the chromosome-level assembly should be treated as preliminary, and work is ongoing to confirm the CC-2931 karyotype. The final assembly consisted of 17 chromosomes and 125 unplaced contigs, with ~98% of sequence placed on chromosomes.

The preliminary chromosome-level assembly was previously used by Chaux-Jukic, et al. (2021) and is accessible at:
https://raba.ibpc.fr/home//Briefcase/Chlamy_genomes
