## Supplemental Figures for "An ancient clade of *Penelope*-like retroelements with permuted domains is present in the green lineage and protists, and dominates many invertebrate genomes"

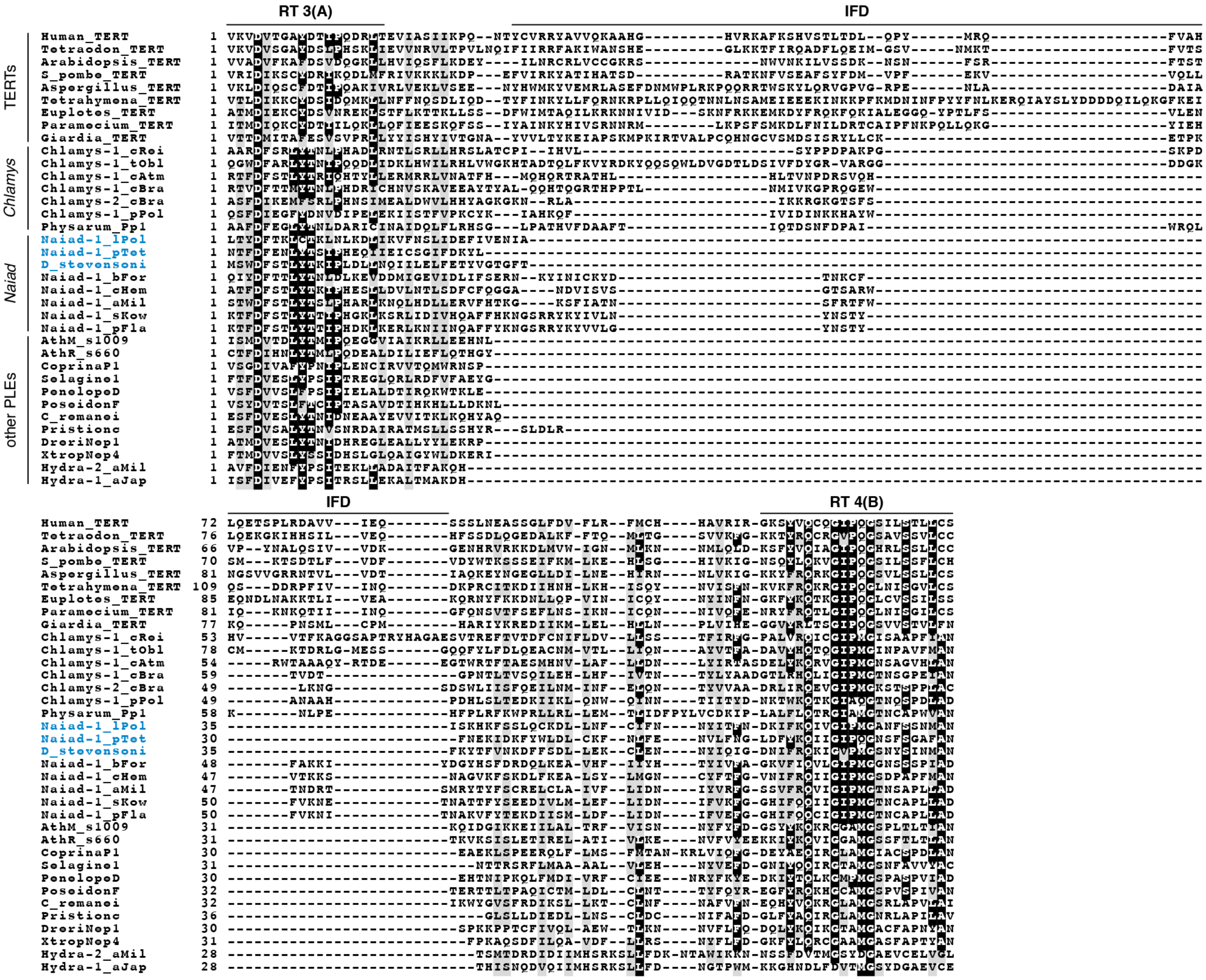


**Figure S1**. Multiple sequence alignment of RT3, IFD and RT4 regions of selected PLEs and TERTs. Examples of *Naiads* lacking the IFD are highlighted in blue text.

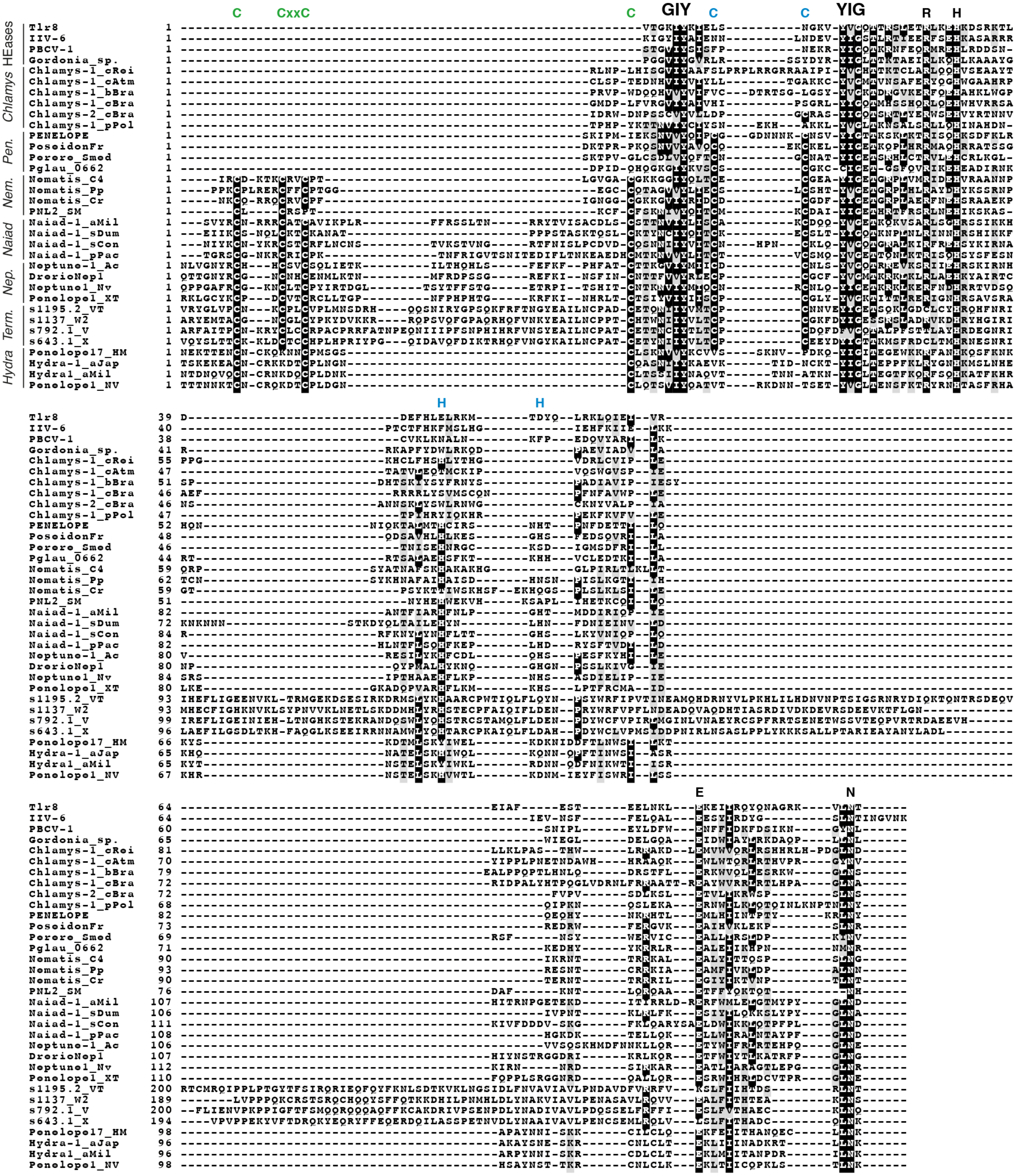


**Figure S2**. Multiple sequence alignment of GIY-YIG ENs across the diversity of PLEs. Select homing ENs that exhibit the highest similarity to PLE ENs are included (see Fig. S8). The conserved aa that form the CCHH Zn-finger in *Penelope/Poseidon*, *Nematis*, *Neptune* and *Naiad* are highlighted in blue, and the aa that form the $CX_{2\text{-}5}CxxC$ Zn-finger in the linker domain of *Nematis*, *Neptune, Naiad* and partially in *Hydra* elements (see Fig. S7) are highlighted in green. *Pen.* = *Penelope/Poseidon*; *Nep.* = *Neptune*, *Nem.* = *Nematis*, *Term.* = *Terminon*.

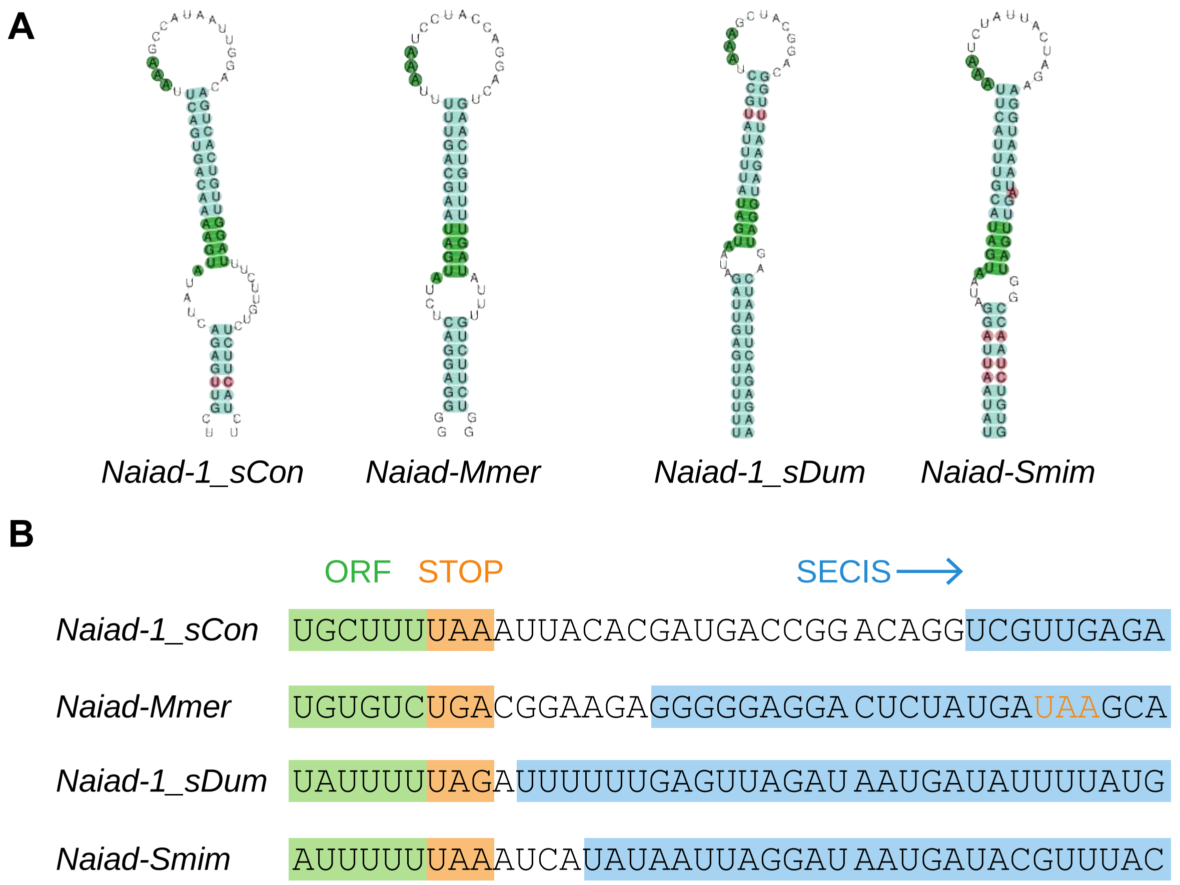


**Figure S3**. SECIS elements in *Naiads* encoding selenoproteins. (**A**) SECIS elements predicted by SECISearch3 (Mariotti et al., 2013). Conserved SECIS features are highlighted in green, standard pairs in cyan and mismatches in red. All SECIS elements were type I (i.e., they consist of two stems and two loops, type II have a third loop). (**B**) Sequence context of inferred stop codons (highlighted in orange) and initial bases of each SECIS element (blue). Note that in *Naiad-Mmer* the stop codon is assumed to be UGA, the first non-UGA stop codon is within the SECIS (UAA, highlighted in orange text).

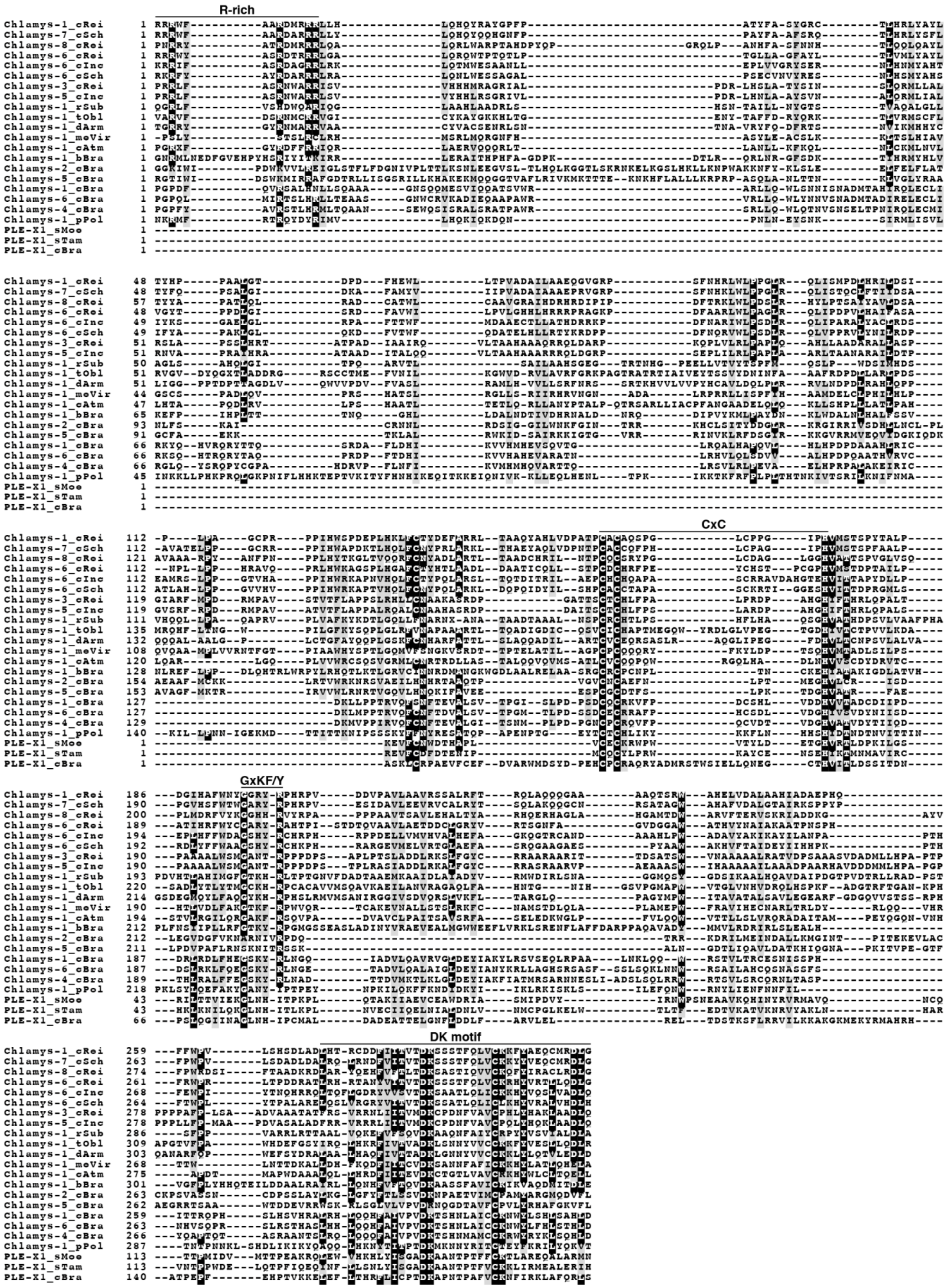


**Figure S4**. Multiple sequence alignment of *Chlamys* elements showing the diagnostic region between EN and the core RT. The R-rich, CxC, GxKF/Y and DK motifs are highlighted. Note that the GxKF/Y and DK motifs are present in PLEs in general, while the R-rich and CxC motifs are unique to *Chlamys* and *Naiads* (see Fig. 3).

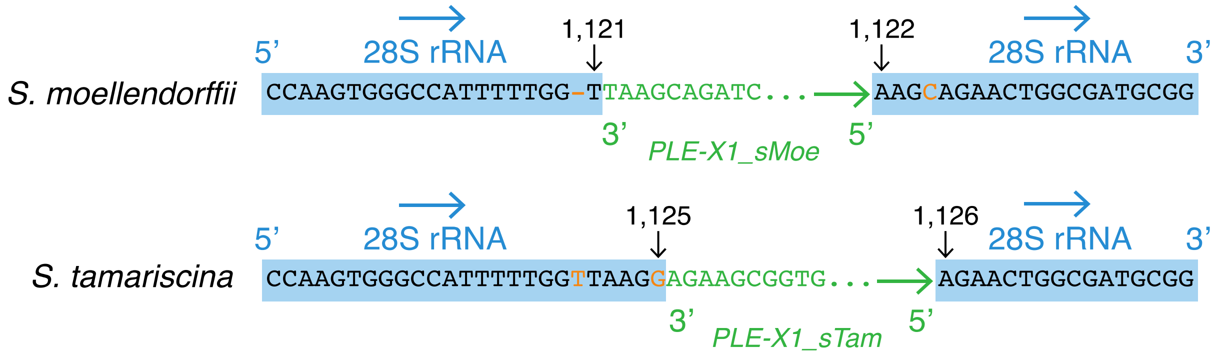


**Figure S5**. Targeted insertion of the 28S ribosomal RNA gene by *Chlamys* elements in *Selaginella*. Nucleotides in blue boxes represent alignment of the 28S rRNA gene between *S. moellendorffii* and *S. tamariscina*, with mismatches highlighted in orange text. Blue arrows represent 5' – 3' orientation. The first 10 bp from the 3' end of each *Chlamys* element are shown in green, with green arrows representing the direction of insertion. The numbered nucleotides flanking the insertions are based on the *S. moellendorffii* 28S rRNA gene sequence (NCBI: XR_002991511.1).

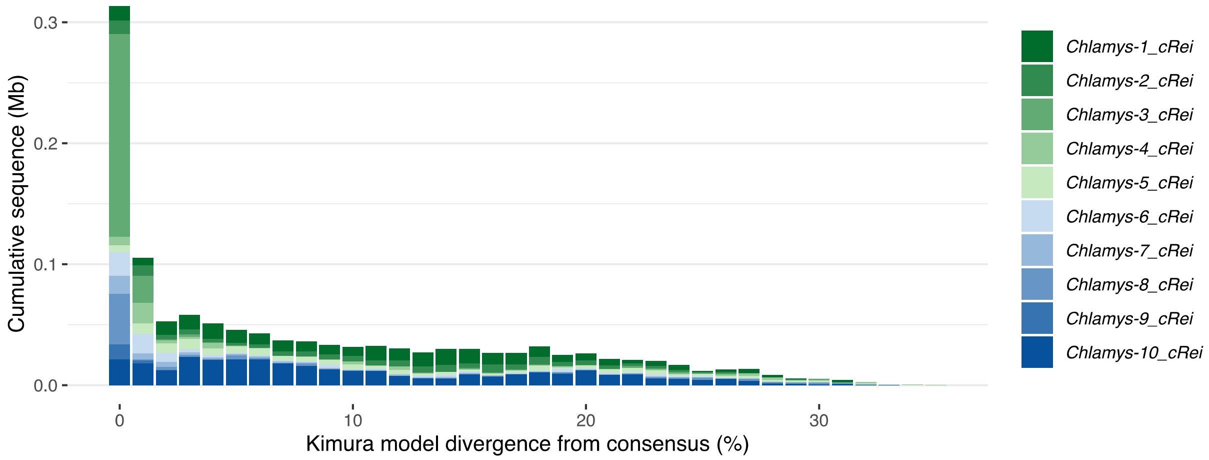


**Figure S6**. Landscape divergence plot for autonomous *Chlamys* elements in *C. reinhardtii*. Plot was generated using the CC-1690 genome assembly (Supplemental Note).

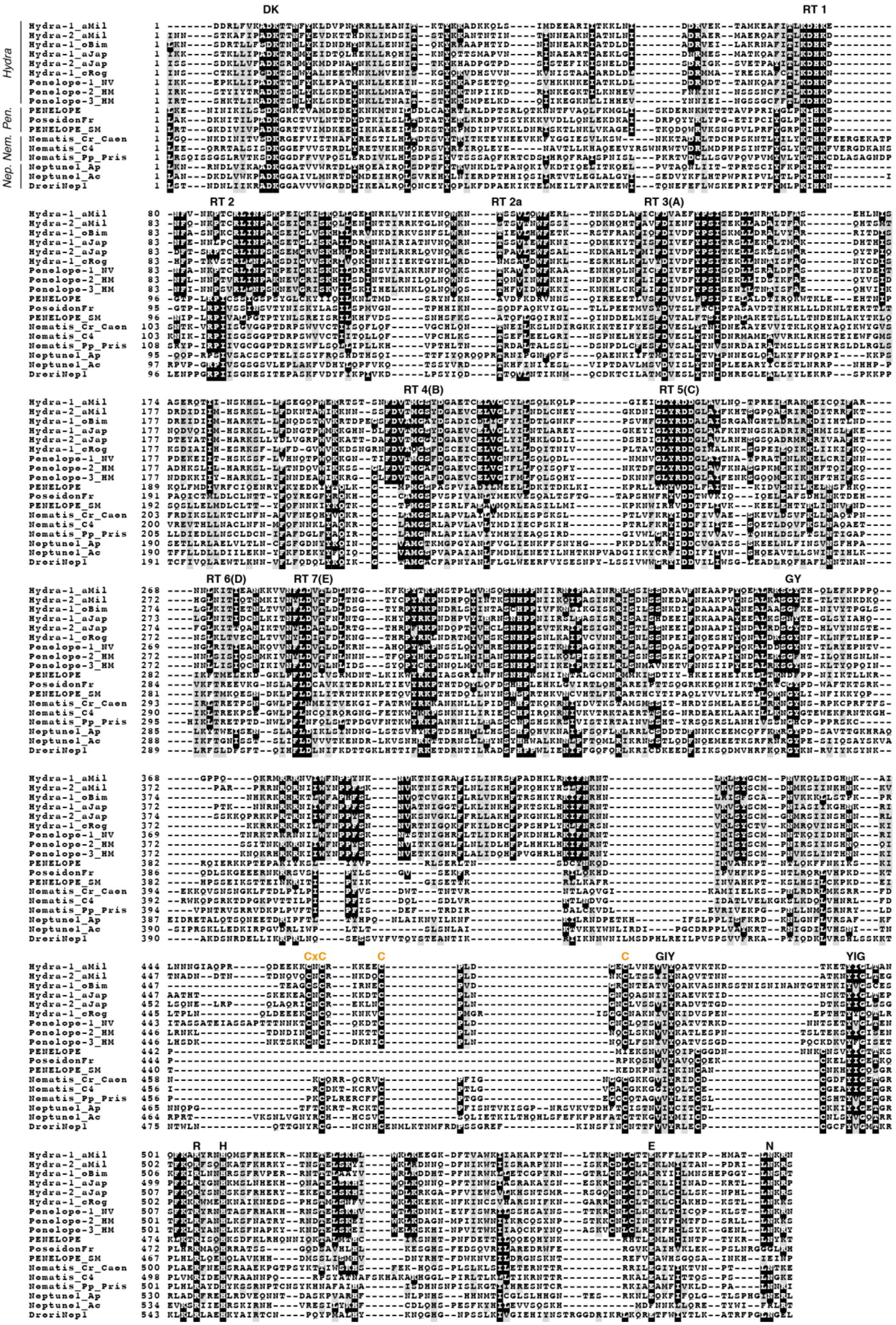


**Figure S7**. Multiple sequence alignment of proteins from the *Hydra* clade and selected canonical PLEs. The conserved cysteines in the modified CxCX$_5$C motif in the EN linker domain are highlighted in orange text. *Pen.* = *Penelope/Poseidon*, *Nem.* = *Nematis*, *Nep.* = *Neptune*.

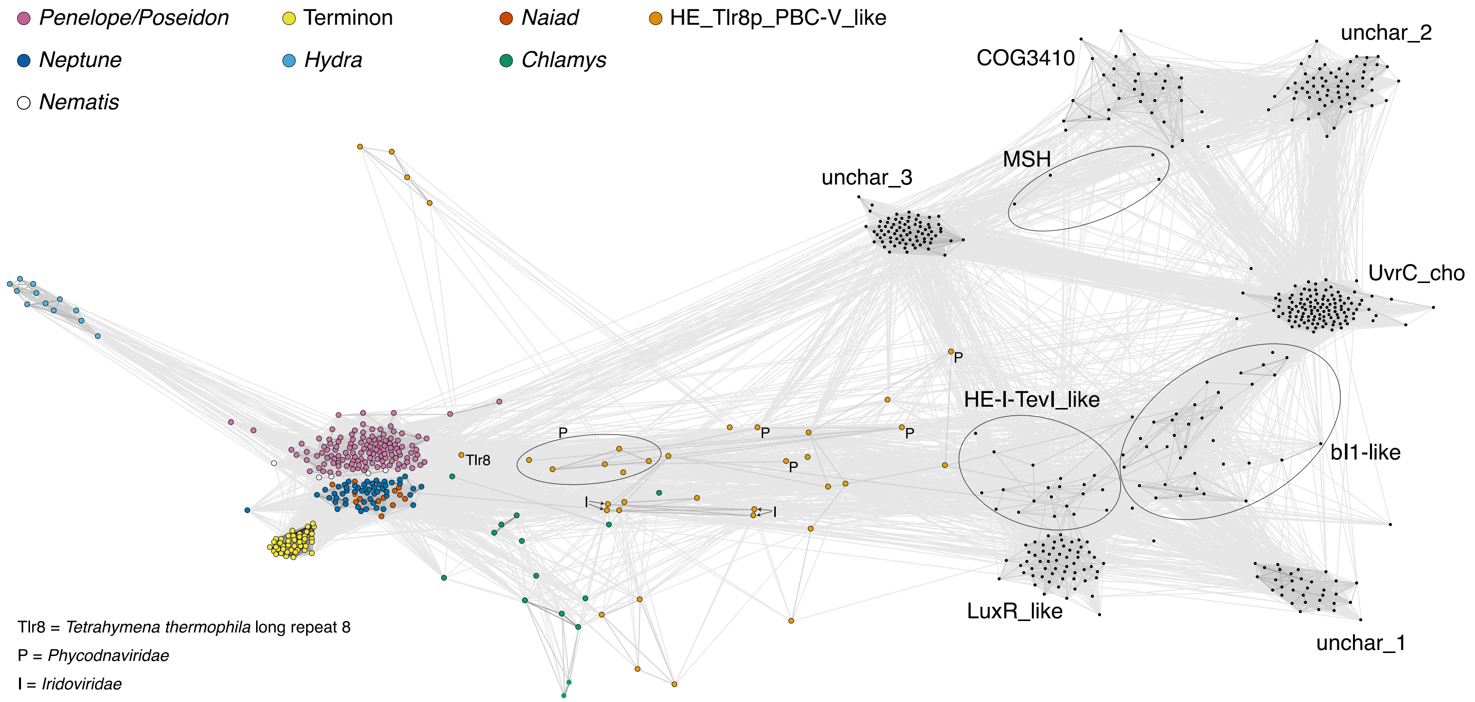


**Figure S8**. CLANS protein clustering of the GIY-YIG EN domain. ENs from PLEs and homing endonucleases of the HE_Tlr8p_PBC-V_like family (cd10443) are colored, and particular HE_Tlr8p_PBC-V_like domains are highlighted. Clusters representing described families from the NCBI GIY-YIG EN superfamily (cd00719) are highlighted.
